# Functional, metabolic and transcriptional maturation of stem cell derived beta cells

**DOI:** 10.1101/2021.03.31.437748

**Authors:** Diego Balboa, Tom Barsby, Väinö Lithovius, Jonna Saarimäki-Vire, Muhmmad Omar-Hmeadi, Oleg Dyachok, Hossam Montaser, Per-Eric Lund, Mingyu Yang, Hazem Ibrahim, Anna Näätänen, Vikash Chandra, Helena Vihinen, Eija Jokitalo, Jouni Kvist, Jarkko Ustinov, Anni I. Nieminen, Emilia Kuuluvainen, Ville Hietakangas, Pekka Katajisto, Joey Lau, Per-Ola Carlsson, Sebastian Barg, Anders Tengholm, Timo Otonkoski

**Author notes:** Equal contribution (Diego Balboa, Tom Barsby, Väinö Lithovius). Corresponding author TO ( / @OtonkoskiT).

## Abstract

Transplantation of pancreatic islet cells derived from human pluripotent stem cells is a promising treatment for diabetes. Despite progress in stem cell-derived islet (SC-islet) generation, detailed characterization of their functional properties has not been conducted. Here, we generated functionally mature SC-islets using an optimized protocol and comprehensively benchmarked them against primary adult islets. Biphasic glucose stimulated insulin secretion developed during in vitro maturation, associated with cytoarchitectural reorganization and increased alpha cells. Electrophysiology and exocytosis of SC-islets were comparable to adult islets. Glucose-responsive insulin secretion was achieved despite differences in glycolytic and mitochondrial glucose metabolism. Single-cell transcriptomics of SC-islets in vitro and throughout 6 months of murine engraftment revealed a continuous maturation trajectory culminating in a transcriptional landscape closely resembling that of primary islets. Our thorough evaluation of SC-islet maturation highlights their advanced degree of functionality and supports their use in further efforts to understand and combat diabetes.

---

The generation of functional pancreatic beta cells from human pluripotent stem cells (hPSCs) is a major goal of stem cell research, aiming to provide a renewable and consistent source of cells for the treatment of diabetes. Stem-cell derived beta cells could solve the limitations of using cadaveric donor islets for transplantation, while serving as a powerful model system to understand the pathogenic mechanisms leading to various forms of diabetes ^1^. Several studies have applied multi-stage *in vitro* differentiation protocols to hPSCs that mimic the sequential inductive signals controlling pancreatic islet development *in vivo*, generating cell clusters (SC-islets) that closely resemble primary islets ^2–9^. Individual studies have reported particular transcriptomic ^7,10^, functional ^3,8,9,11^ and metabolic ^2,12^ aspects of SC-islets. However, studies integrating these aspects with detailed analyses of stimulus-secretion coupling and exocytosis machinery of functional SC-islets have been lacking.

Here, we developed an optimized protocol for the generation of functional SC-islets. We comprehensively compared SC-islets and primary human adult islets to systematically quantify the similarities and differences. During the final, extended maturation stage, the cytoarchitecture of SC-islets was significantly reorganized, and glucose-stimulated insulin secretion matured to a level comparable to primary adult islets. We conducted detailed physiological characterization of the SC-islets, including dynamic insulin secretion assays, respirometry measurements, electrophysiological profiling as well as imaging of Ca^2+^, cAMP and insulin granule exocytosis. These functional assays were further supported by targeted metabolite-tracing studies of glucose flux into the major pathways coupling metabolism to insulin secretion, together with single cell transcriptomic profiling of differentiating endocrine cell populations. Importantly, this multi-pronged approach was conducted during the time-course of SC-islet maturation *in vitro* and following *in vivo* engraftment. Our integrated analyses show that a remarkably high level of beta cell functionality is achieved *in vitro*, even if specific metabolic and transcriptomic differences persist between SC-islet beta cells and primary beta cells.

## Results

### Stem cell derived islets present organotypic cytoarchitecture and function

We devised an optimized differentiation protocol by combining previous advances in the generation of SC-islets ^8,9,13,14^. It employs a maturation medium in the final stage (S7) containing an antiproliferative^15^ aurora kinase inhibitor ZM447439 (ZM), tri-iodothyronine (T3) and N-acetyl cysteine (NAC) (adapted from Patent WO2017222879A1) (Fig.1a). We characterized hESC-derived SC-islet maturation from the beginning of S7 (S7w0) to the end of the sixth week (S7w6) using a series of morphometric, functional, metabolomic and transcriptomic analyses.

**Figure 1.**
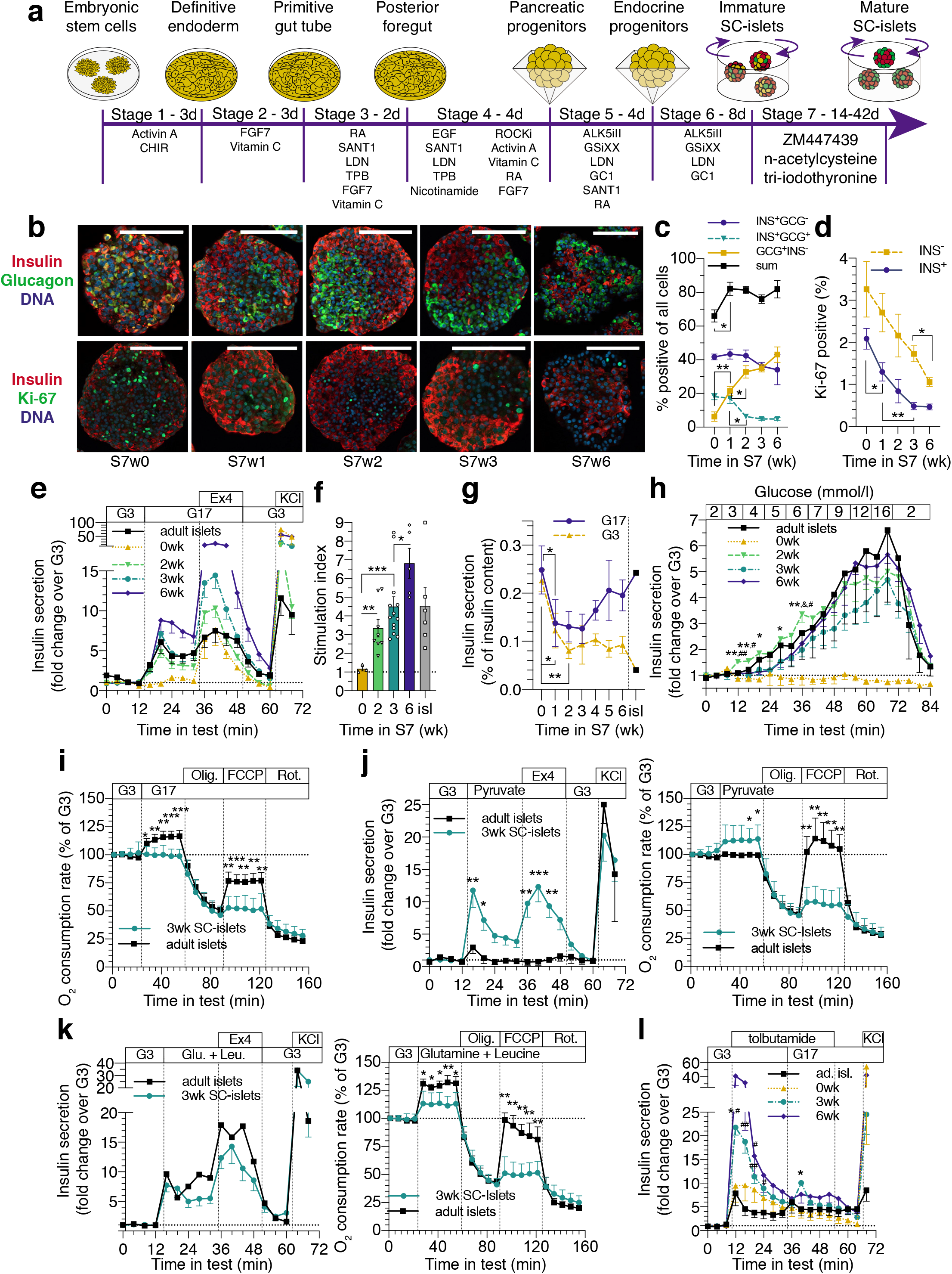
SC-islets achieve cytoarchitecture and insulin secretion characteristics comparable to primary adult islets. **(a)** SC-islet differentiation protocol. Stages 1-4 in monolayer, Stage 5 in microwells, Stages 6-7 in suspension culture. **(b)** Immunohistochemistry of SC-islets during S7 culture, scale bars 100 *μ*m. **(c-d)** Quantification of cell populations during S7 culture from immunohistochemistry: **(c)** INS^+^GCG, INS^-^GCG^+^, INS^+^GCG^+^ and the sum of these populations, % of all cells. n=2-8, Two-way ANOVA, not corrected for multiple comparisons, not assuming the same SD. **(d)** INS^+^Ki67^+^ and INS^-^Ki67^+^ populations, % of the INS^+^ or INS^-^ population, respectively. n=2-8, One-way ANOVA with Welch’s correction, INS^+^ and INS^-^ populations analysed separately. **(e)** Insulin secretion responses to perifusion with 2.8 mM (G3) to 16.8 mM glucose (G17), 50 ng/ml exendin-4 (Ex4) and 30 mM KCl. Normalized to average secretion during G3, the first 16 minutes of the test. n= 3-10. **(f)** Insulin secretion in G17 (average of 16 – 32 min) over secretion in G3 (average of 0 – 12 min) in test (e). Isl=adult islets. One-way ANOVA with Welch’s correction. **(g)** Insulin secretion as percentage of total insulin content in G3 and G17 in 30-minute static incubations. Isl=adult islets. n=3-5 for SC-islets, n=1 for the adult islets. Two-way ANOVA. **(h)** Insulin secretion response to gradual increase in glucose concentration from 2 to 16 mM. Normalized to the average secretion during the first 8 minutes of the test. ‘*’ significance vs. 0wk-, vs. 3wk-, ‘#’ vs. 6wk SC-islets. N= 3-5. Two-way ANOVA, not corrected for multiple comparisons, not assuming the same SD. **(i)** Change in oxygen consumption rate in response to G17, oligomycin (2 *μ*M), FCCP (2 *μ*M) and rotenone (1 *μ*M) in S7w3 SC-islets (n=15) and adult islets (n=5). Student’s unpaired t-test. **(j)** Left: the same test as (e) (n=2-4) and right: the same test as (i) with pyruvate 10mM replacing G17 (n=4-12). Two-tailed Student’s unpaired t-test, with Welch’s correction. **(k)** Left: the same test as (e) (n=1-4) and right: the same test as (i) with glutamine 10mM + leucine 5mM replacing G17 (n=4-11). Two-tailed Student’s unpaired t-test, with Welch’s correction. **(l)** Insulin secretion responses to change from G3 to G17 under the influence of 500 *μ*mol/l tolbutamide (Tolb) in perifusion. Normalized to the average secretion during the first 12 minutes of the test. N=4-5 for SC-islets, N=3 for adult islets. ‘.’ significance vs. 0wk SC-islets, ‘#’ vs. Adult islets. Two-way ANOVA, not corrected for multiple comparisons, not assuming the same SD. All data are presented as mean ± SEM. * p < 0.05, ** p < 0.01, *** p < 0.001

At S7w0, SC-islets presented with ≈40% insulin-positive monohormonal cells (INS^+^), a proportion that remained stable until S7w6, as determined by immunohistochemistry (Fig.1b-c) and flow cytometry (Supp.Fig.1a-b). At S7w0 and S7w1, SC-islets contained ≈20% cells co-expressing insulin and glucagon (INS^+^GCG^+^), a proportion that was reduced to <5% by S7w3. Concomitantly, the number of single-positive GCG^+^ cells increased from ≈5% to ≈40-50% (Fig.1c, Suppl. Fig.1b), consistent with previous studies demonstrating polyhormonal to alpha cell differentiation *in vitro* and *in vivo* ^7,10,16–22^.

While the beta cell numbers remained unchanged, the insulin content of SC-islets increased 6-fold during the first 3 weeks of S7 (Suppl. Fig.1c). Concurrently, SC-islet beta cells progressively acquired dense core insulin granules with ultrastructural morphology resembling those of primary beta cells (Suppl. Fig.1d).

The cytoarchitecture of SC-islets changed significantly during S7 maturation (Fig.1b). A high proportion of INS^+^ cells localized to the SC-islet periphery at the start of S7 (S7w0-w1), but by S7w3, SC-islets were polarized with GCG^+^ and INS^+^ cells clustered at opposing sides. However, by S7w6, the polarization was resolved into a cytoarchitecture with intermingled GCG^+^ and INS^+^ cells (Fig.1b). Quantitatively, this reorganization resulted in an increased number of cell-cell contacts between single-positive GCG^+^ and INS^+^ cells from S7w0 to S7w6 (Suppl. Fig.1e). Importantly, the cytoarchitectural changes during S7 mirror those observed in human pancreatic development ^23–25^.

Functional maturation of developing beta cells is linked to reduced proliferation ^26,27^. SC-beta cell proliferation was reduced by 80% during S7 (from 2.1 to 0.46%) (Fig.1b, d). Non-beta cell proliferation was also reduced (Fig.1d). Critically, the reduced proliferation was dependent on the use of ZM, NAC and T3 within the S7 media (Suppl. Fig.1f), but these agents did not affect SC-islet cell composition (Suppl. Fig.1g).

Adult primary islets are characterized by a tightly controlled, biphasic insulin secretion response to increases in glucose ^28^. We therefore examined dynamic insulin secretion from maturing SC-islets using perifusion assays. At S7w0, high glucose concentrations (16.7 mM) failed to trigger insulin secretion, unless combined with the GLP1 analogue exendin-4, whereas membrane depolarization with high K^+^ triggered a pronounced secretory response. From S7w2, SC-islets showed biphasic glucose-stimulated insulin secretion (GSIS) responses comparable to adult islets, with gradual increases in their stimulation index until S7w6 (Fig.1e, f). This biphasic secretion response was robustly replicated using two additional human iPSC-lines (Suppl. Fig.1h). Of note, the second phase response to high glucose could be sustained continuously for >70 minutes (Suppl. Fig.1i). The acquisition of glucose responsiveness by S7w2 was associated with reduced insulin release in both basal (2.7 mM) and stimulatory glucose concentrations, compared to S7w0. In contrast, the further improvement by S7w6 was due to increased stimulated secretion while low basal secretion was maintained (Fig.1g). The inappropriate basal insulin release at S7w0 was reduced with diazoxide, indicating aberrant stimulus-secretion coupling upstream of K_ATP_-channel closure (Suppl. Fig.1j). Omission of either ZM or NAC from the S7 medium attenuated the GSIS response, while omitting all additives nearly abolished it (Suppl. Fig.1k, l), an effect mostly explained by the higher insulin release in low glucose (Suppl. Fig.1m).

Beta cell functional maturity is reflected in the glucose concentration that triggers insulin secretion ^29^. We assessed the insulin secretion threshold by gradually increasing the glucose concentration in perifusion assays. SC-islets at S7w0 showed no glucose-induced insulin release, whereas at S7w2 they responded at an unphysiologically low glucose concentration of ≈3 mM. In contrast, S7w3 SC-islets displayed the appropriate physiological threshold at ≈5 mM glucose (Fig.1h).

Glucose-induced mitochondrial respiration is another characteristic feature of functional adult islets, which correlates with GSIS ^2,30–32^. We assayed oxygen consumption rates during glucose stimulation (Fig.1i) and observed that glucose increased mitochondrial respiration in primary islets but not in SC-islets, despite comparable insulin secretion dynamics (Fig 1e, 1i). In contrast, SC-islets responded with increased mitochondrial respiration rates (and insulin secretion) to high concentrations of pyruvate, while primary islets remained unresponsive (Fig 1j). Direct stimulation of mitochondrial metabolism using glutamine and leucine triggered comparable insulin release in both primary islets and SC-islets, while the increase in respiration rates was slightly higher for primary islets (Fig 1k). Importantly, basal respiration rates between SC-islets and primary islets were not measurably different under non-stimulatory glucose concentrations (Suppl. Fig.1n). However, with all stimuli, the primary adult islets showed significantly higher mitochondrial spare respiratory capacity (Fig. 1i-k), indicating that mitochondria in the SC islets are less capable to adapt when challenged, and are possibly functionally immature at S7w3.

Finally, to investigate the relative contribution of the triggering and amplifying pathways ^33^ of GSIS, we exposed the SC-islets to tolbutamide, which maximally activates the triggering pathway, followed by high glucose to activate the metabolic amplifying pathways. The S7w3-6 SC-islets demonstrated a stronger insulin secretion response to tolbutamide than adult islets, but a delay of the subsequent glucose-dependent amplification (Fig.1l).

These results demonstrate the generation of SC-islets *in vitro* with biphasic glucose dependent insulin secretion comparable to adult islets and the ability to respond to mitochondrial substrates and incretin receptor stimulation. This acquisition of function correlated with changes in SC-islet architecture and cell composition but not with beta cell mass. However, despite a strong insulin secretion profile, SC-islets showed little glucose-coupled mitochondrial respiration and maintained an inappropriate insulin secretion response to exogenous pyruvate.

### Intact insulin secretory machinery in SC-beta cells

Next, we dissected the stimulus-secretion coupling machinery of the SC-islet beta cells with measurements of ion channel conductance, cytoplasmic Ca^2+^ and cAMP, and exocytosis. Patch-clamp recordings showed that dispersed S7w2 SC-beta cells fired action potentials (Fig.2a) and presente d voltage-dependent Ca^2+^- and Na^+^-currents with voltage dependences similar to those in primary human beta cells (Fig.2b-c). Ca^2+^-current amplitude was similar in both cell types, while Na^+^-currents were about 2-fold larger in SC-beta cells (Fig.2c).

**Figure 2.**
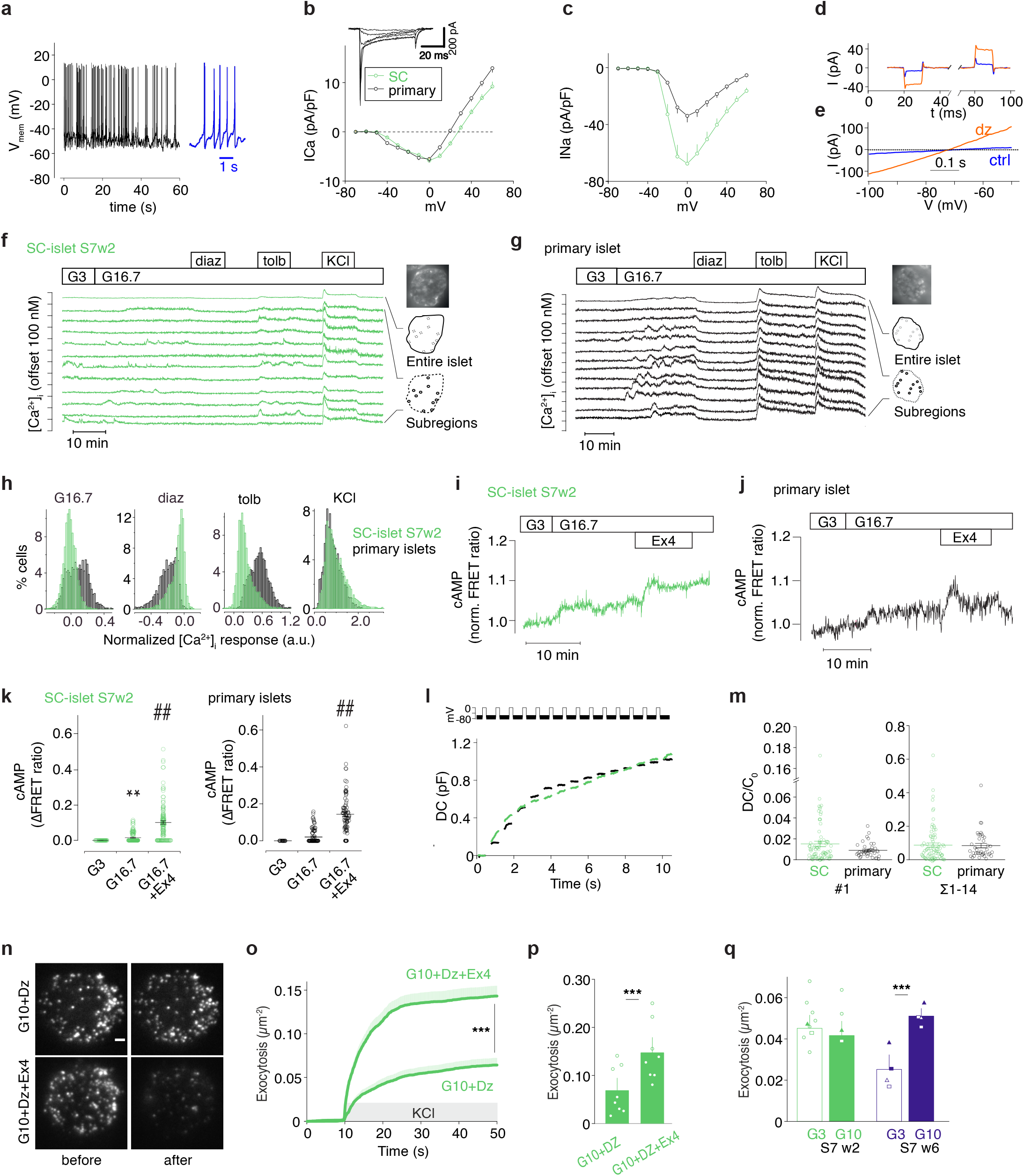
Voltage-dependent ion currets, [Ca2+]i and [cAMP]m signaling and exocytosis in SC-derived and in adult islet cells. **(a)** Example membrane potential recording in beta cells of dispersed SC-islets; 10 mM glucose. **(b-c)** Current (I)-voltage (V) relationship of inward currents in beta cells of dispersed SC-islets (green n = 80 cells, 8 preparations) or primary islets (black, n=39 cells, 4 donors). Inset shows family of voltage-clamp currents in SC-beta cells (−40 and +10 mV from a holding potential of −70mV). Average Ca2+ current during 5–45ms of the depolarization (b) and peak Na+ current during the first 5ms of the depolarization p = 0.002, two-tailed t-test. (c); currents are normalized to cell capacitance (pF). For SC-beta cells, half-maximal Ca2+-current activation was reached at −29.0±0.9 mV (n=64) and half-maximal Na+-current activation was at −22.5±0.4 mV (n=75). **(d-e)** Current responses to step-depolarizations (d; +/-10 mV around −70 mV, black) or voltage ramps (e; −100 to −50 mV at 100 mV/s) in ctrl (blue) or in presence of diazoxide (200 *μ*M, orange). **(f-g)** [Ca2+]i recordings from 2wk SC-islets (f) and primary islets (g) exposed to 3 (G3) and 16.7 mM glucose (G16.7), 250 *μ*M diazoxide, 1 mM tolbutamide and 30 mM K+. The uppermost trace shows a quantification from an entire islet and the traces below are representative examples from cell-sized regions of interest. **(h)** Histograms showing the changes of [Ca2+]i in response to various treatments normalized to the levels at G3 in cells from SC-islets (green; n=5254) and primary islets (black, n=3550). **(i-j)** Representative [cAMP]pm recordings from cells in intact SC- (i) and primary (j) islets stimulated with G16.7 and 10 nM exendin-4. **(k)** The effects of G16.7 and 10 nM exendin-4 from experiments in (i) and (j). n=119 cells from 6 independent experiments with SC-islets and 81 cells from 3 primary islet preparations. **, P<0.01 vs 3 mM glucose; ##, P<0.01 vs 16.7 mM glucose, Student’s paired t-test. **(l)** Cell capacitance increase (ΔCm) during a train of 14×200-ms depolarizations from −70mV to 0mV in SC-beta cells (green) and primary beta cells (black). **(m)** Average change in membrane capacitance, normalized to initial cell capacitance (ΔC/C0), during the 1st depolarization (#1), and total increase during the train (Σ1-14) for SC-beta (n = 80 cells, 8 preparations) and primary beta cells (n = 39 cells, 4 preparations). Dots represent individual cells and lines the mean values. **(n)** TIRF micrographs of SC-beta cells expressing the granule marker NPY-tdmOrange2 in absence (top) or presence of exendin-4 (Ex4, bottom), and before (left) and after (right) stimulation with elevated K+; bath solution contained 10 mM glucose and diazoxide. Scale bar 2 *μ*m. **(o)** Time course of average cumulative number of high K+-evoked exocytotic events normalized to cell area in experiments as in (n), for control (68 cells), and Ex4 (71cells). p = 4*10-8, two-tailed t-test. Shaded areas indicate SEM. **(p)** Total K+-depolarization-induced exocytosis in (o). **(q)** Spontaneous exocytosis (normal K+, no diazoxide) during a 3-minute observation period after >20 min pre-incubation at G3 or G10. Fusion events were quantified in SC-beta cells at S7w2 (13 cells at G3 and 12 at G10) and at S7w6 (40 cells at G3 and 41 at G10) and normalized to the cell area. In p-q, dots represent averages for individual SC-islet batches. All data presented as mean ± SEM unless otherwise indicated.

K_ATP_-channel dependent K^+^-conductance of S7w2 SC-beta cells was quantified using symmetric voltage-steps (Fig.2d) or ramps (Fig.2e). In 3 mM glucose, the membrane conductance was on average 53 ± 4 pS/pF (n=50 cells), and increased in presence of the K_ATP_-channel opener diazoxide in 49/50 cells to 273 ± 30 pS/pF (n=50 cells). Both values are similar to those previously reported for human beta cells ^34^.

The cytoplasmic Ca^2+^ concentration ([Ca^2+^]_i_) was recorded in individual cells within intact SC-islets and primary islets. In S7w2 SC-islets, a subset of cells showed [Ca^2+^]_i_ oscillations in low glucose with little change in response to high glucose (Fig.2f). However, many cells showed low and stable [Ca^2+^]_i_ at low glucose, with increased and often oscillatory [Ca^2+^]_i_ in high glucose (Fig.2f). Primary islets also behaved heterogeneously (Fig.2g), but a higher proportion of cells responded to elevated glucose (Fig.2h). K_ATP_-channel opening with diazoxide reduced, and closure with tolbutamide increased [Ca^2+^]_i_ in both SC-islets and primary islets, with more pronounced responses in the primary islets (Fig.2f-h). Depolarization with 30 mM K^+^ increased [Ca^2+^]_i_ in all cells, and with similar magnitude in SC-islets and primary islets (Fig. 2f-h). [Ca^2+^]_i_ responses in SC-islets were similar at the beginning and end of S7 (Suppl.Fig.2a,b), and therefore cannot explain the improved glucose response after prolonged culture (Fig.1e-h). However, among the glucose-responsive cells, basal [Ca^2+^]_i_ decreased from 47±0.4 to 24±0.2% of the K^+^-stimulated level and the increase induced by glucose stimulation improved from 5.2±0.1% at S7w0 (n=1091 cells) to 20.8±0.4% at S7w8 (n=2270 cells), consistent with the observed reduction of basal secretion and improved stimulatory index.

In the presence of low glucose, tolbutamide increased [Ca^2+^]_i_ in both primary and SC-islets, but again to a higher degree in primary islet cells (Suppl.Fig.2c-e). High glucose in the continued presence of tolbutamide caused slight [Ca^2+^] _i_ increase in SC-islets and a decrease in primary islet cells. Despite this lowered [Ca^2+^]_i_, glucose amplified secretion under these conditions (Fig.1l). Activation of GLP-1 receptors with exendin-4 did not alter [Ca^2+^]_i_ in primary islets and had only a weak tendency to increase [Ca^2+^]_i_ in SC-islets (Suppl. Fig.2c-e).

The sub-membrane cAMP concentration ([cAMP]_m_), a modulator of insulin exocytosis, was recorded in single cells within intact SC and primary islets. High glucose induced a small, and exendin-4 a much more pronounced increase in [cAMP]_m_ in both preparations (Fig 2i-k), indicating that cAMP signalling in SC-islets closely resembles that in primary human islets.

Single-cell exocytosis was measured as membrane capacitance changes using patch clamp. A train of depolarizations (14 x 200 ms) resulted in identical capacitance increases (ΔC) of 0.087 ± 0.012 fF/pF (n=80) in S7w2 beta cells and 0.084 ± 0.013 fF/pF (n=39) in primary beta cells (Fig.2l-m). Cell size, as assessed by the initial cell capacitance (Cm_0_), was slightly larger for SC-cells than for primary beta cells (by 27 %, p=0.0001, unpaired t-test) (Supp.Fig.2f).

Docking and exocytosis of insulin granules at the plasma membrane were studied by TIRF microscopy. Expression of the granule marker protein NPY-tdmOrange2 for 26-48 h in S7w1 cells resulted in a punctate granule staining (Fig.2n), and depolarization with elevated K^+^ (in the presence of diazoxide) released 0.063 ± 0.008 granules μm^-2^ (n=68 cells, Fig.2o). Exocytosis proceeded initially with a burst (4 × 10^-3^ gr μm^-2^ s^-1^, <10s) and decreased later to <0.7 × 10^-3^ gr μm^-2^ s^-1^. When exendin-4 was added, we consistently observed more than a doubling of K^+^-stimulated exocytosis (0.14 ± 0.01 granules μm^-2^, n=71 cells, Fig.2o-p). All exocytosis values are similar to those reported by us in primary beta cells ^35^.

Notably, in S7w1, spontaneous exocytosis (no diazoxide) was similar in 3 mM and 10 mM glucose was not different from exocytosis at low glucose (0.045 ± 0.004 gr μm^-2^, n=13 vs. 0.041 ± 0.002 gr μm^-2^, n=12; Fig.2q). In contrast, at S7w6, basal exocytosis in 3 mM glucose was lower (0.025 ± 0.002 gr μm^-2^, n=40) and doubled when glucose was raised to 10 mM (0.051 ± 0.003 gr μm^-2^, n=41; Fig.2q), which is consistent with the improved glucose regulation in this preparation (Fig.1g).

The density of docked granules (visible in TIRF-M) was ~ 0.6 gr/μm^2^ in S7w2 cells (Supp.Fig.2g-k), which is identical to our reported values for primary beta cells ^35^. Treatment with exendin-4 slightly increased docked granules when exocytosis was prevented with diazoxide (Supp.Fig.2g).

In summary, these analyses showed that SC-beta cells are equipped with the necessary ion channels, exocytosis components and intracellular signalling machinery required for fine-tuned regulation of insulin secretion.

### Glucose tracing analysis of SC-islets reveals remodelling, but not full maturation, of glucose metabolism

To further probe the discrepancy between SC-islet functionality and low respiratory coupling we investigated how glucose metabolism differed between primary islets and SC-islets. We performed metabolite tracing analyses using uniformly labelled [U-^13^C6]-glucose comparing S7w0, w3 and w6 SC-islets together with primary adult islets, under low (3 mM) and high labelled glucose (17 mM) conditions (Fig.3a).

**Figure 3.**
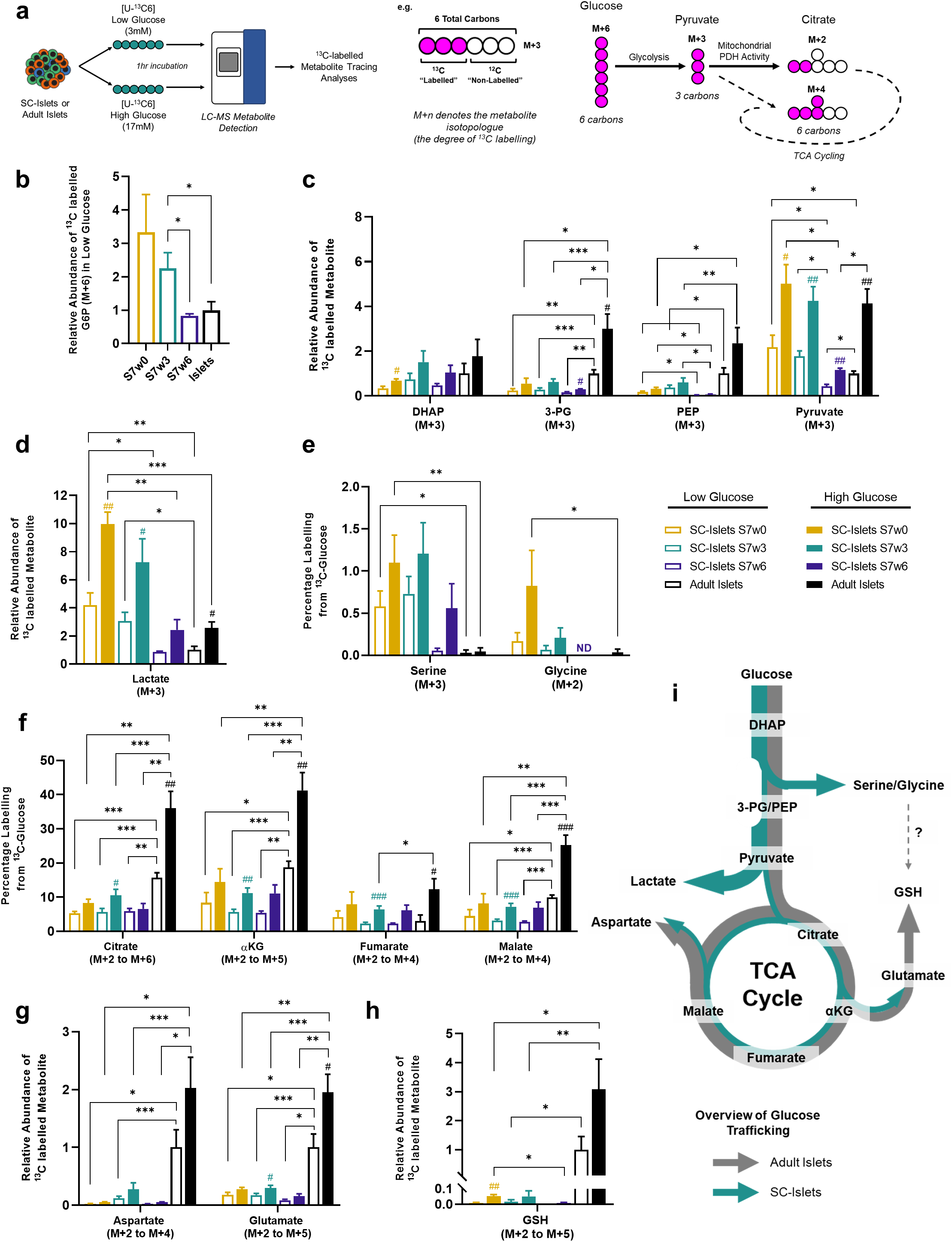
Metabolic tracing analysis of maturing SC-islets. **(a)** (left) Overview of experimental setup. SC-islets or adult islets were exposed to low (3mM) or high (17mM) concentrations of uniformly labelled [U-13C6] glucose for 1 hour before metabolite extraction and liquid-chromatography mass spectrometry (LC-MS) detection. (right) An example of isotopologue nomenclature and glucose-derived labelling of downstream metabolites. **(b)** The relative abundance of M+6 labelled Glucose-6-Phosphate under low glucose treatment of adult islets and SC-islets from week 0 to week 6 of maturation. **(c)** The relative abundances of fully labelled glycolytic intermediates in SC-islets and adult islets following low and high labelled glucose treatment. **(d)** The M+3 lactate content of adult islets and SC-islets detected over the time course of maturation, following low and high labelled glucose treatment. **(e)** The percentage of total serine and glycine labelled from 13C-glucose following low and high glucose treatment of SC- and adult islets. **(f)** The combined percentage of labelled TCA metabolites (M+2 to fully-labelled M+n) from SC- and adult islets after low and high labelled glucose treatment. **(g)** The combined abundance of aspartate (M+2 to M+4) and glutamate (M+2 to M+5) isotopologues in SC-islets (w0 to w6) under low and high glucose concentrations, relative to adult islets. **(h)** The combined relative abundance of M+2 to M+5 glutathione isotopologues under low and high labelled glucose concentrations in adult islets and SC-islets. **(i)** Schematic overview of active glucose metabolic pathways in SC-islets and adult islets. Arrow thickness denotes the extent of glucose-derived carbons entering the pathway. Error bars ± SEM with statistical significance determined by two-tailed t-tests with Welch’s correction. ‘#’ symbols indicate internal significance from low to high glucose labelling, ‘*’ symbols denote significance between SC-islet timepoints or adult islet samples at each glucose concentration. #,*p < 0.05, ##,**p < 0.01, ###,***p < 0.001. SC-islets S7w0 (n=4), S7w3 (n=12-13), S7w6 (n=3), adult islets (n=6).

Beta cell glucose-sensing is in part mediated by the hexokinase step of glycolysis, through the phosphorylation of imported glucose into glucose-6-phosphate (G6P) by glucokinase (GCK) ^31^. Over the course of SC-islet maturation, we detected a reduction in the relative abundance of labelled G6P produced at low glucose (Fig.3b), suggesting a tighter control of glucose phosphorylation upon maturation. This is consistent with the observed reduced insulin secretion (Fig.1e-g) and exocytosis (Fig.2q) at low glucose in more mature SC-islets. Concurrently, residual labelled glucose levels at low glucose conditions were higher in primary islets than SC-islets (Supp.Fig.3a).

We observed reduced labelled and total levels of 3-phosphoglycerate (3-PG) and phosphoenolpyruvate (PEP) in SC-islets compared to primary islets, despite comparable levels of labelled dihydroxyacetone phosphate (DHAP) in low and high glucose conditions (Fig.3c, Supp. Fig.3c). These results are consistent with a proposed glycolytic “bottleneck” due to reduced GAPDH activity ^12^.

A significant decrease in the production of labelled lactate was a strong characteristic of SC-islet maturation during S7 (Fig.3d). This correlates strongly with the tighter control of glucose phosphorylation described above, and is in agreement with studies that link lactate overproduction to reduced GSIS ^36,37^. The diversion of 3-PG into de novo serine and glycine biosynthesis, which is low in primary islets but significantly higher in less mature SC-islets, is another possible avenue of aberrant glucose metabolism (Fig.3e).

SC-islets showed increased labelled glucose incorporation into the core tricarboxylic acid (TCA) cycle metabolites citrate, alpha-ketoglutarate (αKG), fumarate and malate upon stimulation with high glucose, but the response was clearly lower than in primary islets (Fig.3f). The overall lower glucose-derived carbon flux into the TCA cycle in SC-islets, in contrast with robust GSIS, suggests a lower threshold of mitochondrial activation necessary for glucose stimulated insulin secretion. We inferred flux through the TCA cycle by tracing the degree of ^13^C-glucose derived carbon incorporation into each TCA metabolite, which increases with every rotation of the TCA cycle, generating distinct isotopologues (e.g. M+2, M+4). Primary islets showed enhanced cataplerotic TCA cycling for citrate, αKG, fumarate and malate compared to SC-islets (Supp.Fig.3c-f), as well as enhanced flux through anaplerotic reactions, via M+3 oxaloacetate from M+3 pyruvate, that resulted in a high proportion of M+3 malate and fumarate isotopologues (Supp.Fig.3e-f). Even though SC-islets presented reduced cycling of glucose-derived carbons (both for anaplerotic and cataplerotic reactions) than adult islets, the extent of M+2 and M+3 isotopologue labelling was comparable, and neither TCA cycle direction seemed aberrantly biased (Fig.3f, Supp.Fig.3c-f).

Aspartate and glutamate are key components of the malate-aspartate redox shuttle that transfer reduced NADH moieties into the mitochondrial matrix, a key component of beta cell metabolism that supports glucose stimulated insulin secretion ^38^. Primary islets used a significantly higher proportion of glucose-derived carbons (in both low and high glucose) to generate these amino acids than SC-islets (Fig.3g). Also, a significantly higher amount of labelled glutathione (GSH) was produced by primary islets than SC-islets (Fig.3h), which is mirrored by their higher production of the GSH precursor glutamate. In contrast, the synthesis of labelled serine and glycine (another GSH component) was barely detectable in primary islets as seen above (Fig.3e). Of note, primary islets maintained higher total (i.e. labelled and unlabelled) levels of reduced and oxidized forms of glutathione (GSH and GSSG) and the electron carriers NAD and NADP (Supp. Fig.3g).

Thus, primary human islets and SC-islets not only differ in their core TCA cycle turnover and respiration rates under glucose stimulation, but also in the production of TCA-derived metabolites. Despite the differences in both glycolytic and mitochondrial glucose metabolism (Fig.3i), SC-islets do display dynamic glucose-sensitive insulin secretion responses, suggesting that the acquisition of the missing metabolic couplings would further enhance and fine-tune their functionality.

### SC-islets control the glycaemia of mice *in vivo*

To investigate *in vivo* functional potential of the immature (S7w0) and more mature (S7w3) SC-islets, we implanted them under the kidney capsule of non-diabetic mice^2,9,10,39–41^ (Fig4a). Circulating human C-peptide was detectable at 1-month post-engraftment in all engrafted mice. However, mice engrafted with S7w3 SC-islets demonstrated 2-fold higher human C-peptide levels at 2 and 3 months than S7w0 engrafted mice (Fig.4b). Correspondingly, blood glucose levels at 3 months were lower in S7w3 SC-islet engrafted animals (Fig.4c) and reached the human glycaemic set point (4.5 mM) by 3 months post-engraftment, as reported in primary islet engraftment studies^42^. Glucose tolerance tests showed regulated insulin secretion in response to glucose injection in mice carrying both types of grafts (Fig.4d), but the glucose clearance was more rapid in S7w3 engrafted mice (Fig.4e,f). Next, we tested whether the S7w3 SC-islet grafts could sustain normoglycaemia after streptozotocin (STZ) induced loss of endogenous murine beta cells. Glucose tolerance tests before and after STZ treatment showed that both control and STZ-treated animals presented glucose-regulated C-peptide secretion (Fig.4g). Despite C-peptide levels being lower in the STZ group, the glucose levels were similarly controlled in both groups (Fig.4h). The proportions of INS^+^ and GCG^+^ cells were not affected by the STZ treatment (Fig.4i,j). After removal of the engrafted kidney, the blood glucose levels increased sharply (Fig.4k), demonstrating that the engrafted SC-islets were actively controlling the glycaemia of the diabetic mice. Extended *in vitro* culture in S7 conditions thus confers SC-islets a degree of maturation that results in improved functionality upon engraftment *in vivo*.

**Figure 4.**
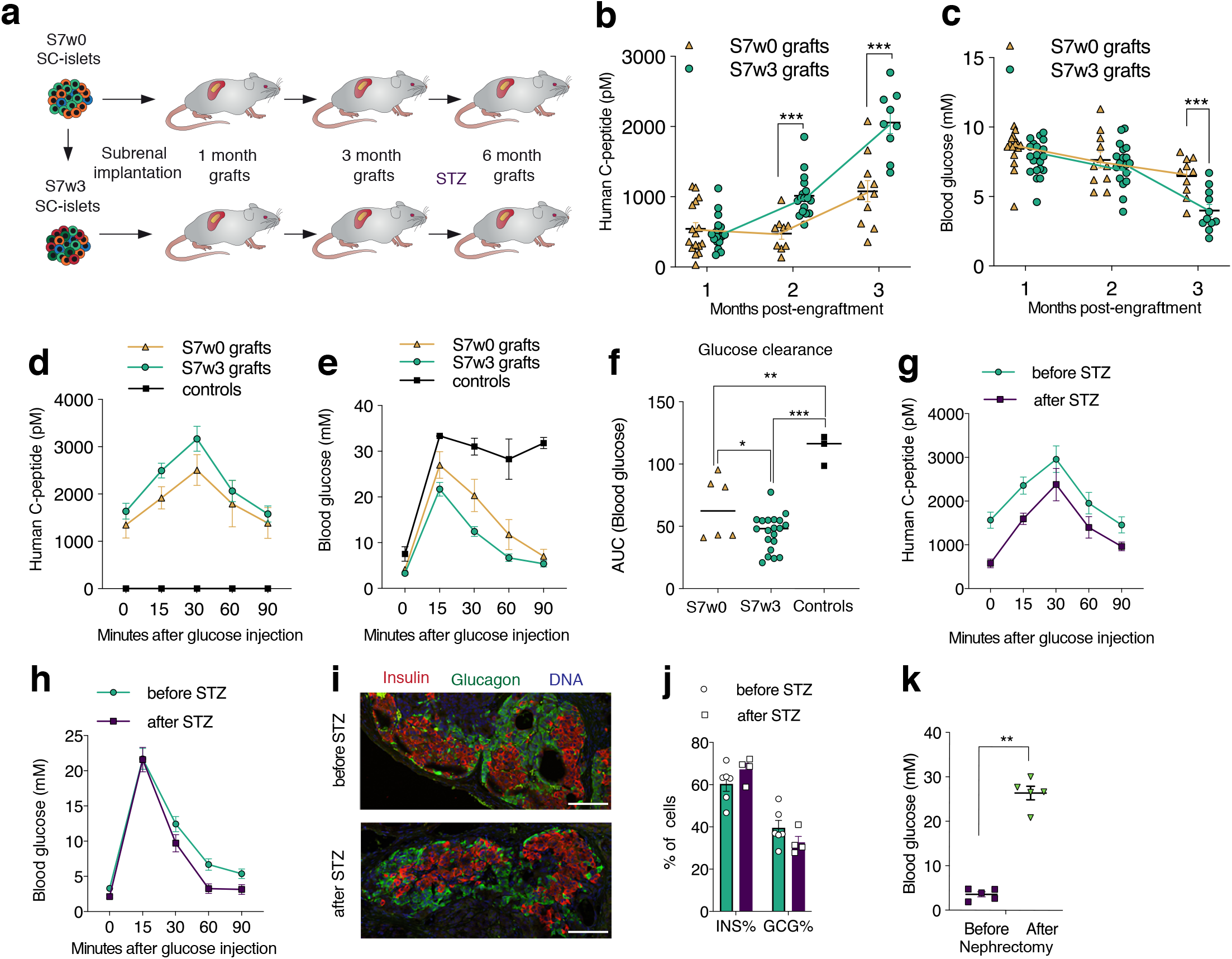
Engrafted SC-islets provide effective glycemic control in mice. **(a)** Schematic representation of the in vivo experiments. **(b-c)** Measurements form random-fed mice engrafted with S7w0 or S7w3 SC-islets, (b) Human specific C-peptide levels (pM) and (c) Mouse blood glucose levels (mM) (S7w0: n=16 in 1m, 9-11 in 2m and 10-11 in 3m; S7w3: n=18-20 in 1m, 17-19 in 2m and 9-11 in 3m). Individual data points presented, Multiple t-test. **(d-e)** Human C-peptide (c) and blood glucose (d) levels measured during an intraperitoneal glucose tolerance test (IPGTT) at three months post-engraftment. Mice were injected with 3 mg/kg glucose after a 6 h fast. (S7w0: n=6, S7w3: n=20, non-implanted controls n=3). **(f)** Glucose clearance quantified from IPGTT blood glucose values in (d) as area under the curve (AUC). One-way ANOVA and Tukey’s multiple comparisons test. **(g-h), (h-i)** Human C-peptide levels (pM) **(g)** and blood glucose levels (mM) **(h)** in an IPGTT similar to the one in **(d-e)** in mice at 4 months post-engraftment (before streptozotocin (STZ) injection, n=10) and at 5 months post-engraftment (after STZ injection, n=9). **(i)** Immunohistochemistry in the grafts before and after STZ injection. Scale bar 100*μ*m. **(j)** Quantification of INS+ and GCG+ populations before (n=6) and after (n=4) STZ injection, % of all cells positive for either INS or GCG. Dots represent graft-wide data from individual grafts. **(k)** Blood glucose levels from engrafted, random-fed mice before and after nephrectomy at 5-6 months post-engraftment. The mice were injected with streptozotocin (STZ) at 4 months post-engraftment. (n=4) Mann-Whitney U-test. All data represent mean ±SEM, unless indicated otherwise * p < 0.05, ** p < 0.01, *** p < 0.001.

### SC-islets transcriptionally mature *in vitro* and *in vivo*

To further elucidate the molecular mechanisms behind the observed *in vitro* and *in vivo* maturation, we performed single cell RNA sequencing at four timepoints of SC-islet *in vitro* differentiation (S5, S7w0, S7w3 and S7w6), as well as SC-islet grafts retrieved at 1-, 3- and 6 months post-engraftment (Fig. 5a). After integration with previously published datasets ^43,44^, we generated a dataset containing 42 261 pancreatic endocrine cells that clustered in different populations according to time of origin and cell identity (Fig.5b,c; Supp.Fig.4a-d; Supp.Table1,2). These populations reconstructed a differentiation continuum, from multipotent pancreatic progenitors expressing *PDX1, NKX6-1* and *NEUROG3;* through intermediate differentiating stages, into beta (44%), alpha (33%), delta and gamma cells (Fig.5c-d, Supp.Table1-2, Supp.Fig.4e-i). We detected an increase in alpha cells and a decrease in polyhormonal cells during extended S7 culture (Suppl. Fig.4h), in line with the immunohistochemistry and flow cytometry results.

**Figure 5.**
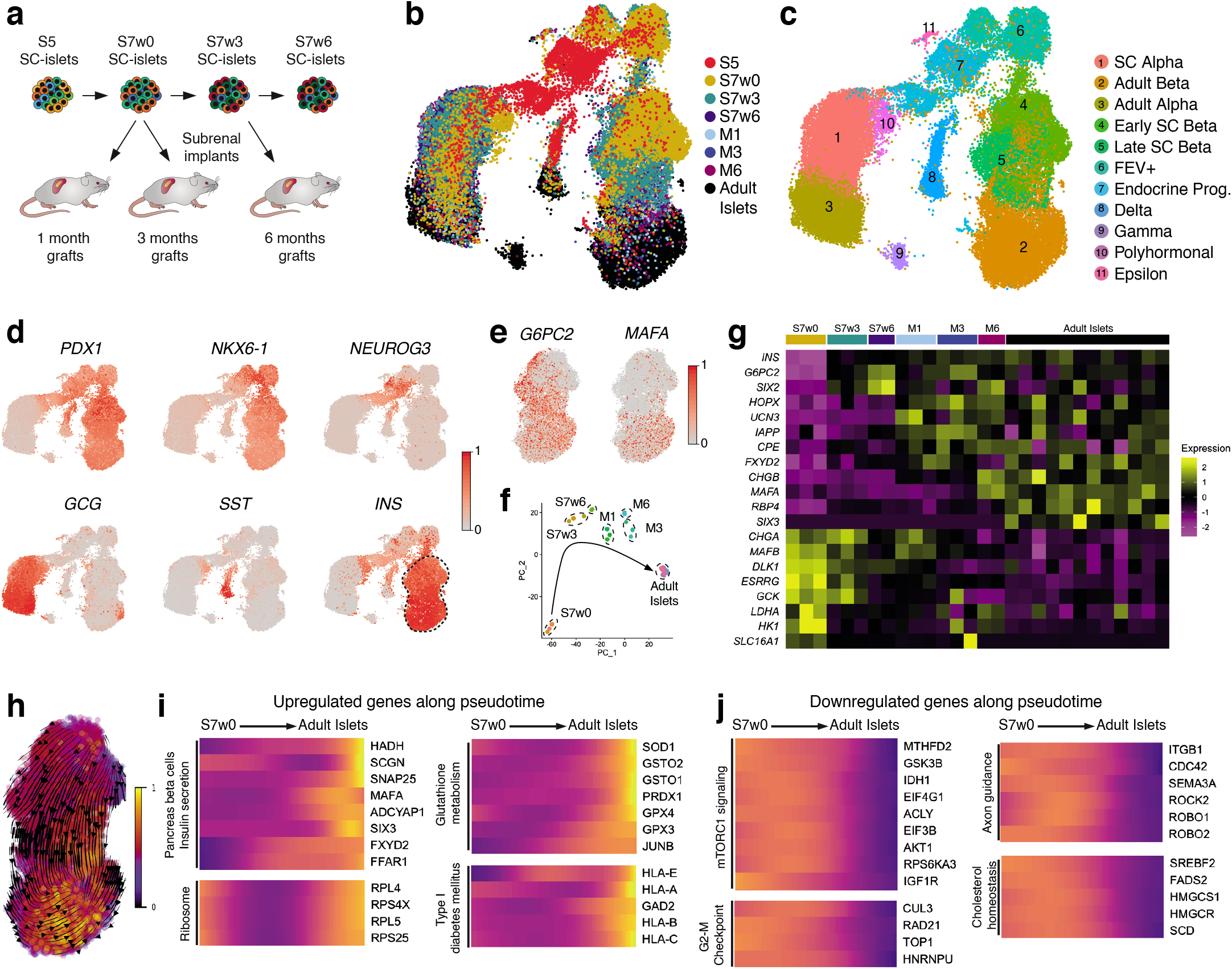
Single cell transcriptomic profiling of stem cell derived islet cells. **(a)** Experimental outline for scRNAseq transcriptomic profiling of SC-islets at the end of in vitro culture stages 5 (S5) and 6 (S7w0) and at week 3 (S7w3) and week 6 of S7 culture (S7w6), together with grafts retrieved after 1 (M1), 3 (M3) and 6 months (M6) post-engrafment. **(b)** Uniform Manifold Approximation and Projection (UMAP)-base embedding projection of an integrated dataset of a total of 46 261 SC-islet and adult human islet cells (from Krentz et al. 2018 and Xin et al. 2018), coloured by time of origin. **(c)** Clustering of the dataset in (b) cells into different cell types. **(d)** Relative expression of marker genes for pancreatic progenitor cells (PDX1, NKX6-1, NEUROG3) and alpha-(GCG), delta-(SST) and beta-(INS) cells. **(e)** UMAP projection of the beta cell population indicating the relative expression of mature beta cell markers G6PC2 and MAFA. **(f)** Principal component analysis of the independent scRNAseq dataset samples. (S7w0: n=3; S7w3: n=3; S7w6: n=2; M1: n=3; M3: n=3; M6: n=2;Adult: n=12). **(g)** Average expression of beta cell maturation marker genes across samples from different time or origin. **(h)** Pseudotemporal ordering of beta cells in (e) based on RNA velocities, colours represent pseudotime. **(i-j)** Relative expression levels of example genes that are upregulated (i) or downregulated (j) along pseudotime.

We focused on the beta cell subpopulations to determine transcriptional changes associated with functional maturation (Fig.5d-e, Suppl. Fig.4k). Principal component and correlation analyses indicated that S7w3 and S7w6 beta cells *in vitro* were transcriptionally more similar to grafted and adult beta cells than to S7w0 beta cells (Fig.5f, Suppl. Fig.4j). We examined the average expression of known mature beta cell marker genes across each individual sample ^7,10,45–48^ (Fig.5g). *INS* gene expression increased over time as did *G6PC2* and *SIX2*, which were already highly expressed during *in vitro* culture. Other mature beta cell marker genes such as *HOPX, UCN3, IAPP, CPE* and *FXYD2* were markedly upregulated upon engraftment. *CHGB* and *MAFA* expression was sharply upregulated at 6 months post-engraftment (M6), suggesting the need of extended *in vivo* maturation for the regulation of these genes, whereas *RBP4* and *SIX3* were mainly expressed in adult beta cells (Fig.5g). Interestingly, while *CHGB* levels increased with time, the expression of its paralog gene, the pan endocrine marker *CHGA*, decreased upon maturation, together with transcription factor *MAFB*. The expression of glucose-phosphorylating enzymes glucokinase *GCK* and hexokinase *HK1*, a beta cell disallowed gene ^49^, was reduced from S7w3 onwards, while their counteracting enzyme, phosphatase *G6PC2*, was upregulated over time. This strengthens the notion of tightening glycolytic flux control upon beta cell maturation, in accordance with the functional and metabolic findings (Fig.1h, Fig.3b). The expression of the disallowed genes *SLC16A1* (a pyruvate/lactate transporter) and lactate dehydrogenase *LDHA*^50,51^ peaked at S7w0 and was thereafter downregulated, consistent with the lowered lactate production during *in vitro* SC-islet maturation (Fig.3d), but incompletely inhibiting the functional response to pyruvate retained by *in vitro* SC-islets (Fig.1j).

RNA velocity estimation and pseudotemporal ordering of the beta cells yielded a directional differentiation trajectory that enabled us to investigate genes differentially regulated upon beta cell maturation (Fig.5h). The expression of genes associated with pancreatic beta cells and insulin secretion increased with pseudotime, together with ribosome, HLA-genes, and glutathione metabolism genes (Fig.5h, Supp. Fig.4l-m, Supp.Table.3). Conversely, the expression of genes related to mTORC1 and MAPK signalling, cholesterol homeostasis, mitosis and MYC targets (i.e. proliferation) decreased with pseudotime. Also, genes associated with axon guidance (*ROBO1, ROBO2*) and adherence junctions were downregulated, indicating changes in cell migration, adhesion and cytoskeletal properties, which is consistent with the observed cytoarchitectural changes upon maturation (Fig.5j, Supp.Fig.5l-n, Supp.Table3).

We then investigated the transcriptional changes associated with *in vitro* maturation by comparing S7w0, S7w3 and S7w6 SC-beta cells (Fig.6a-b, Supp.Table.4). Genes associated with beta cell maturation and function were upregulated from S7w0 to S7w6 (*INS, IAPP, CHGB, G6PC2, FXYD2, TFF3, WNT4, GABRA2*), together with serine biosynthesis genes (*PSAT1, SHMT2, PHGDH*) and early response genes (*FOS, JUN, JUNB*). Gene sets for beta cell ion channels (*KCNJ11, CACNA1D*), unfolded protein response (*ERO1A, WFS1*), and TGF-beta signalling (*BMP2, TGFB2, TGFBR3, ID1*) were enriched in the S7w6 upregulated genes (Fig.6c, Supp.Fig.5a,b). Downregulation of genes related to MYC targets, cholesterol homeostasis and disallowed genes was again observed in more mature S7w6 beta cells (Fig.6b-c, Suppl.Fig.5b). The expression of oxidative phosphorylation-related genes is markedly higher in S7w0 beta cells, lowest in S7w3 and then increases in S7w6 and *in vivo* stages (Fig.6c, Suppl. Fig.4n, Suppl. Fig.5b). Transcription factor *MAFB* ^52^ and its targets *ACVR1C, DLK1* and *CRYBA2* were also downregulated (Fig.6f, Supp.Fig.5a-b, Supp.Table5).

**Figure 6.**
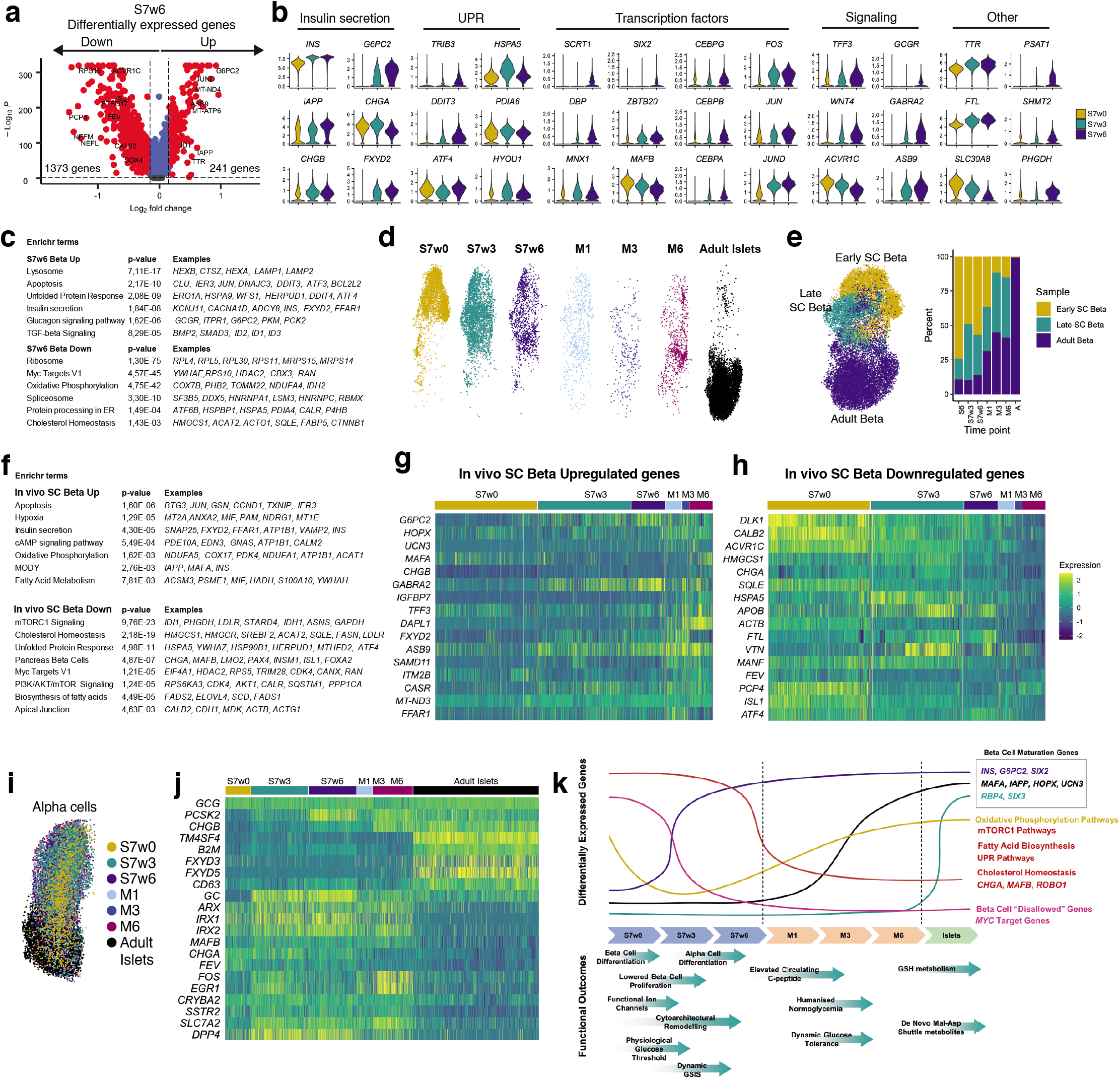
Transcriptional maturation of stem cell derived beta cells. **(a)** Differentially expressed genes between S7w6 and S7w0-3 SC-beta cells. **(b)** Selected genes from different functional categories that are differentially expressed across in vitro SC-beta cell maturation. **(c)** Gene sets enriched in the S7w6 SC-beta cells upregulated and downregulated genes. **(d)** Distribution of the beta cells from different time of origin within the beta cell cluster. **(e)** Clustering of beta cells according to their transcriptional similarity into early, late and adult beta like profiles and the fractional contribution to each category from beta cells of different time of origin. **(f)** Gene sets enriched in the in vivo implanted vs in vitro SC-beta cells. **(g-h)** Expression of genes upregulated and down-regulated in the in vivo SC-beta cells. **(i)** Distribution of alpha cells from different time of origin within the alpha cell cluster. **(j)** Expression of marker genes for each alpha cell subpopulation by their time or origin. **(k)** Summary of functional and transcriptomic features of SC-islet maturation in vitro and in vivo.

The unfolded protein response (UPR) pathway, which senses ER-stress levels, was elevated in S7w3-6 compared to S7w0, with upregulation of *TRIB3, DDIT3, ATF4, HSPA5* in parallel with increased *INS* levels (Fig.6b,c). Genes involved in cell cycle inhibition (*TP53, CEBP* transcription factor family^53^, and *SCRT1* ^54,55^), were upregulated in S7w6 beta cells, while MYC target genes were downregulated (Fig.6b, Supp.Fig.5a). All together, these transcriptional changes further indicate increased ER-stress in S7w6 beta cells, together with reduced proliferation, both characteristic features of mature beta cells.

Beta cell differentiation is not a synchronous and homogeneous process, as evidenced by the co-clustering of beta cell signatures from different time points along the pseudotemporal trajectory (Fig.6d-e). To reduce the noise introduced by heterogeneous populations, SC-derived beta cells were unbiasedly clustered by transcriptionally similarity, rather than by time of origin, into SC-early, SC-late and Adult beta categories (Fig.6e). The proportion of beta cells in the SC-late and Adult beta categories increased with the progression of time *in vitro* and *in vivo* (Fig.6c). Adult beta category cells upregulated genes related to mature insulin granule content and secretion (*PCSK1, CPE, CHGB, ABCC8*), beta cell maturation (*MAFA, GDF15*) and mitochondrial oxidative phosphorylation (Suppl.Table.6).

During *in vivo* maturation after SC-islet engraftment (Fig.4, Fig.6f-h, Suppl. Fig.5a, Supp.Table.5), SC-beta cells further upregulated genes related to beta cell maturation and function, insulin secretion, cAMP signalling pathway, oxidative phosphorylation and fatty acid metabolism. Grafted SC-beta cells downregulated genes associated with mTORC1 signalling, cholesterol homeostasis, biosynthesis of fatty acids and apical junction components, together with beta cell developmental transcription factors (Fig.6f).

SC-alpha cell populations also displayed transcriptional changes during their maturation *in vitro* and *in vivo*, upregulating genes related to alpha cell glucagon granules (*GCG, PCSK2, CHGB, GC*) (Fig.6i-j). Consistent with the increased differentiation of SC-alpha cells during S7 and the augmented alpha-beta cell interaction due to cytoarchitectural reorganization (Fig.1b-c, Supp. Fig.1b,e) S7w6 beta cells presented increased expression of the glucagon receptor (GCGR) (Fig.6b, Supp. Fig.5d, Supp.Table.7) alluding to increased interplay of these endocrine cell types during maturation.

## Discussion

We describe the generation of SC-islets with insulin secretory function closely matching that of primary adult human islets. SC-islets developed robust biphasic insulin responses to glucose together with a physiological glucose threshold. Compared to other recently published protocols ^2–4,6,12,56^, this protocol leads to similar or improved insulin secretory function without the need of purification or reaggregation steps. The functional maturation observed over time in S7 was not driven by the generation of more beta cells but instead correlated with decreased beta cell proliferation, increased alpha cell fraction and progressive rearrangement of SC-islet cytoarchitecture ^27,57,58^.

Electrophysiological analyses in SC-beta cells confirmed exocytosis, K_ATP_ and calcium channel activity similar to primary islets, which also was supported by [Ca^2+^] _i_ recordings. SC-islet cells still presented less pronounced glucose-induced [Ca^2+^]_i_ changes than primary islets, likely reflecting a limited glucose depolarizing effect, since high K^+^ triggered a much stronger [Ca^2+^]_i_ elevation and insulin release. However, GSIS was comparable to that in adult islets, which may reflect the involvement of amplifying pathways controlling the magnitude of insulin secretion. Indeed, cAMP signalling, which enhances Ca^2+^-triggered exocytosis, did not differ between SC-islets and primary islets. Also, transcriptomic profiling showed that cAMP signalling downstream of the glucagon receptor was improved in S7w6 SC-beta cells, in parallel with SC-alpha cell differentiation and enhanced alpha-beta cell interaction upon SC-islet cell reorganization ^58^. Notably, SC-islets presented an enhanced insulin secretory response to GLP-1 receptor activation and membrane depolarization compared to primary islets.

Over the course of *in vitro* maturation, the glucose-sensitivity of insulin granule exocytosis increased, resulting in higher release in elevated glucose and lower release in basal glucose. The latter was paralleled by a reduction of basal [Ca^2+^]_i_ and is in agreement with the increased glucose threshold of insulin secretion in perifusion assays. These findings may be explained by the tightened control of glucose phosphorylation during maturation, as indicated by the decrease in labelled G6P at basal glucose, downregulation of *GCK* and *HK1* genes, and upregulation of *G6PC2* phosphatase.

Diversion of glucose carbons into lactate and serine production likely contributes to reduced late glycolysis metabolite levels and mitochondrial metabolism stimulation. Similarly, we observed a glycolytic restriction in SC-islets^12^ beyond GAPDH which may explain the low mitochondrial respiration response to high glucose, but allowing pyruvate responsiveness. Lactate production and pyruvate response are mediated by the disallowed genes *LDHA* and *SLC16A1* respectively, which in the adult beta cell are selectively repressed to prevent the metabolism of non-glucose carbon sources that would abnormally elicit insulin secretion^59^ and to restrict the anaerobic respiration of glucose^36^. Importantly, *LDHA* and *SLC16A1* showed a pattern of downregulation during the early stages of *in vitro* SC-beta cell maturation.

Beta cell maturation has also been linked with enhanced mitochondrial oxidative phosphorylation, ostensibly powered by increased TCA cycle metabolite production under glucose stimulation^60,61^. As has been previously reported in other models of SC-beta maturation ^2^, we observe mitochondrial gene upregulation from S7w3 onwards. Of note, the highest expression of oxidative phosphorylation genes occurred in immature S7w0 SC-beta cells, probably related to their active proliferation status, which is in line with findings in immature mouse beta cells ^62^. However, the increased expression of oxidative phosphorylation genes was not mirrored by adult level respiration rates and TCA cycle turnover.

Pathways cycling TCA metabolites between the cytosol and the mitochondria, such as the PEP cycle^12,63^, the pyruvate-isocitrate cycle^64^, the pyruvate-malate cycle^61^, and the malate-aspartate redox shuttle^65^ (involving glutamate) contribute to the coupling of beta cell glucose metabolism with insulin secretion^38^. In our metabolic dataset we did detect high levels of anaplerotic TCA metabolite labelling, which would fuel such cycles. Cytosolic redox reactions have also been implicated in glucose stimulated insulin release^64,66^. We observed upregulation of many glutathione (GSH)-related synthesis and redox genes during SC-beta maturation, while detecting much higher levels of de novo GSH production in adult islets using targeted metabolomic analysis.

Despite these differences in glucose metabolism, SC-islets acquired a robust insulin secretory function, suggesting that secondary pathways derived from TCA metabolites (such as glutamate, aspartate, and GSH) may have a beneficial effect on glucose sensitivity without stark increases in aerobic metabolism^38,67,68^.

The transcriptomic profiling of SC-islets upon *in vitro* and *in vivo* maturation revealed significant expression changes in important pathways and processes involved in islet cell maturation. We detected downregulation of mTORC1 signaling-related genes, in particular after engraftment. This is consistent with its reported role in postnatal mouse beta cell maturation and islet development ^69^, where a switch from mTORC1 to AMPK signaling mediates a shift to oxidative metabolism and functional maturation ^70,71^. During SC-islet maturation we also found downregulation of genes related to cholesterol homeostasis and the biosynthesis of fatty acids, processes that play a key role in beta cell function ^31,38,72^.

Chronic endoplasmic reticulum stress (ER-stress) induced by the high rate of insulin biosynthesis is a hallmark of post-mitotic beta cells ^73,74^. We found increased expression of ER-stress related genes in mature *in vitro* SC-beta cells, correlating with their higher expression of insulin and reduced proliferation. Among these upregulated genes we detected *TRIB3*, an AKT-signalling inhibitor that has been linked to beta cell cycle arrest^75^ and is induced by chronic ER-stress associated with insulin biosynthesis. *SCRT1*, a gene involved in repressing beta cell proliferation and the regulation of insulin secretion ^54,55^, was also upregulated.

In summary, we present a protocol for the reliable generation of physiologically relevant SC-islets with cytoarchitecture and functionality comparable to adult primary islets. Moreover, the multi-faceted analysis that we present here constitutes a comprehensive effort to thoroughly benchmark maturing SC-islets against human primary adult islets, considered the “gold standard” in the field. This in-depth parallel characterization of functional, metabolic and transcriptomic aspects of SC-beta cells through *in vitro* and *in vivo* maturation constitutes a valuable resource to further understand the mechanisms of human islet development and functional maturation (summarized in Fig.6k). It also demonstrates the progress in achieving functionality similar to adult islets, while indicating the directions to enhance transcriptional and metabolic hallmarks. The combination of these integrated analyses with refined differentiation protocols will guide the generation of further improved SC-islets for the modelling of beta cell dysfunction, drug-screening purposes and cell replacement approaches, expanding our understanding of the disease mechanisms and therapeutic possibilities to combat diabetes.

## Methods

### *In vitro* culture and differentiation of hPSCs

Human embryonic stem cell line H1 (WA01, WiCell) was used in the majority of the study. In supplementary figure 1h, iPSC lines HEL24.3 ^76^ and HEL113.5-corrected ^77^ were used as well. The iPSCs were generated and used according to the approval of the coordinating ethics committee of the Helsinki and Uusimaa Hospital District (no. 423/13/03/00/08). The hPSCs were cultured on Matrigel (Corning, #354277) coated plates in Essential 8 (E8) medium (Thermo Fisher, A1517001) and passaged using EDTA. To prepare the differentiation experiments, near-confluent plates of stem cells were dissociated using EDTA and seeded on new Matrigel coated plates in E8 supplemented with 5-10 μM Rho-Associated kinase inhibitor (ROCKi, Y-27632, Selleckchem S1049) at a density of ≈0.22 million cells/cm^2^ to achieve confluent plates. To start the differentiations, the medium was changed to D0 medium 24h post-seeding. The differentiation was carried out using a 7-stage protocol combined from key publications^8,9,13,14^ and the patent WO2017222879A1. Complete media formulations are available in the Supplementary Table 8. On the 3^rd^ day of S4 culture, the planar pancreatic epithelium was dissociated using 6-10 min TrypLE treatment (Thermo Fisher, 12563029) and seeded to microwells (6-well AggreWell 400 plates, Stem Cell Technologies) at a density of 800-1000 cells per microwell using the manufacturer’s recommended protocol. On the 1^st^ day of S6 culture the SC-islets were transferred from the microwells to suspension culture on ultra-low attachment plates (Corning, CLS3471) placed on rotator spinning at 95 RPM. Media changes were performed daily until the 1^st^ day of S6 culture and every two to three days thereafter. SC-islet differentiations were carried out by 5 different researchers and all produced SC-islet batches of similar functionality.

### *In vitro* culture of primary adult islets

Primary islets were provided by the Nordic Network for Islet Transplantation (Uppsala University) and University of Alberta IsletCore (Canada). They were maintained in CMRL1066 supplemented with 10% FBS, 20 mM HEPES (Gibco, 15630-056), 2 mM Glutamax and 100 IU/ml penicillin and 100 μg/ml streptomycin on ultra-low attachment plates. Islet donor characteristics are listed in Supplementary Table 9.

### Flow cytometry

Stage 7 SC-islets were dissociated with TrypLE for 5–10 min at 37°C water bath and resuspended in 5% FBS-containing PBS. Cells were fixed and permeabilized using Cytofix/Cytoperm (554714, BD Biosciences) for 20 minutes. Primary antibodies were incubated overnight at 4°C and secondary antibodies for 30 min in RT in Perm/Wash buffer (554714, BD Biosciences) containing 5% FBS. The cells were run on FACSCalibur cytometer (BD Biosciences) and analysed with FlowJo v10 (BD Biosciences). Antibodies are listed in the Supplementary Table 10.

### Immunohistochemistry and image analysis

Samples of S7 SC-islets were fixed for 2h and samples of explanted SC-islet grafts fixed overnight in 4%PFA and embedded in paraffin. Five μm-sections were deparaffinized and subjected to HIER in 0.1 mmol/l citrate buffer. The slides were blocked with UV-block (Thermo Scientific, TA-125-PBQ), and incubated with primary antibodies diluted in 0.1% tween-20 overnight in +4°C. Secondary antibodies were diluted similarly and incubated for 1h in RT. Antibodies are listed in the Supplementary Table 10. The slides were then imaged with Zeiss AxioImager using Apotome II with the same exposure and export setting used on all slides of each immunostaining. Immunostainings were analysed using CellProfiler ^78^ 4.0 with pipelines similar to the ones previously reported ^77^. In brief, the nuclei were identified first and expanded along the intensity gradient of the cytoplasmic staining. Nuclei were assigned to the cytoplasm if 25% of the nuclear perimeter was overlapping with the corresponding cytoplasm. For the neighbour-to-neighbour analyses, each identified insulin+ nucleus was expanded by 35 pixels, and each overlapping insulin+ or glucagon+ or hormone negative nucleus was identified as its neighbour. The same pipeline settings were used on all images of each immunostaining, and the thresholds were set using the robust background algorithm.

### *In vitro* tests of insulin secretion

Static tests of insulin secretion were carried out in 1.5ml tubes. 20-30 SC-islets were hand-picked and equilibrated in Krebs-Ringer-buffer (KRB) with 2.8 mM glucose (G3) for 90 minutes, and then subjected to sequential 30-min incubations of G3, 16.8 mM glucose (G17) and 30 mM KCl or G3, G3 + 100 μM diazoxide, G17 + 100 μM diazoxide and 30 mM KCl in KRB. Dynamic tests of insulin secretion were carried out using a perifusion apparatus (Brandel Suprafusion SF-06, MD, USA) with a flow rate of 0.25 ml/min, and sampling every 4 minutes. 50 hand-picked SC-islets were used for each channel. The SC-islets were perifused with KRB and sample collection was started after 90 minutes of equilibration in G3. Six different test protocols were used with conditions labelled in the corresponding figures. Samples from each collected fraction were analysed using insulin ELISA (Mercodia, Sweden). After static and dynamic tests of insulin secretion, the SC-islets were collected and the insulin and DNA contents were analysed.

### Respirometry

Respiration rates of SC-islets and primary islets were measured using a Seahorse Bioscience XFe96 Extracellular Flux Analyzer. S7w3 SC-islets and primary islets were loaded into Matrigel -coated Seahorse XF96 cell culture microplates (20-25 per well) and attached to the surface overnight in S7 growth media. Prior to the assay, the medium was exchanged into KRB containing 3mM glucose, and the islets allowed to equilibrate for 60-90 minutes at 37°C. Oxygen consumption rates (OCR) were measured over 150 minutes. Basal OCR was calculated prior to the sequential addition of a stimulatory nutrient (17mM glucose, 10mM pyruvate, or 10mM glutamine and 5mM leucine), an inhibitor of ATP-synthetase activity (2μM oligomycin), a mitochondrial uncoupling agent (2μM Carbonyl cyanide-4-(trifluoromethoxy)phenylhydrazone, FCCP), and finally an inhibitor of Complex I of the electron transport chain (1μM rotenone) in KRB. Respiration rates were normalized to the basal OCR prior to nutrient or small molecule addition. To compare the absolute level of OCR between SC-islets and adult islets, raw OCR values were normalized to the DNA content of each well.

### Transmission electron microscopy

Samples from SC-islets and human islets were chemically fixed with 2.5% glutaraldehyde (EM-grade, Sigma-Aldrich) in 0.1 M sodium cacodylate buffer, pH 7.4, supplemented with 2 mM calcium chloride at RT, for 2 h. After washing, the specimens were osmicated in the same buffer with 1% non-reduced osmium tetroxide on ice, for 1 h. Specimens were then washed and dehydrated in increasing concentration of ethanol and acetone, prior to gradual embedding into Epon (TAAB 812). After polymerization over 18 h at 60°C, a pyramid was trimmed on the location of the embedded cells. Ultrathin, 60-nm sections were cut using an ultramicrotome (Leica ultracut UCT, Wetzlar, Germany), picked on a Pioloform coated single slot grids and post-stained with uranyl acetate and lead citrate. Micrographs were acquired with a Hitachi HT7800 microscope (Hitachi High-Technologies Corp., Tokyo, Japan) operated at 100 kV using a Rio9 CMOS-camera (AMETEK Gatan Inc., Pleasanton, CA). One or two SC-islets were randomly chosen for examination and 11-12 SC-islet beta cells were selected for imaging based on the characteristics features of beta-like granules. **Electrophysiology.** SC-islets were dispersed into single cells by gentle pipetting in cell dissociation buffer (Thermo Fisher Scientific) supplemented with trypsin (0.005%, Life Technologies). The dispersed cells were then washed and plated in serum-containing medium onto 22-mm polylysine-coated coverslips, allowed to settle overnight, and then transduced with adenovirus coding for EGFP under control of the RIP2 promoter to identify beta cells.

Standard whole-cell voltage clamp and capacitance recordings were performed using an EPC-9 patch amplifier (HEKA Electronics, Lambrecht/Pfalz, Germany) and PatchMaster software. Voltage-dependent currents were investigated using an IV-protocol, in which the membrane was depolarized from −70 mV to +80 mV (10 mV steps) during 50 ms each. Currents were compensated for capacitive transients and linear leak using a *P*/4 protocol. Exocytosis was detected as changes in cell capacitance using the lock-in module of Patchmaster (30 mV peak-to-peak with a frequency of 1 kHz).

Patch electrodes were made from borosilicate glass capillaries coated with Sylgard close to the tips and fire-polished. The pipette resistance ranged between 2 and 4 MΩ when filled with the intracellular solution containing (in mM) 125 Cs-glutamate, 10CsCl, 10NaCl, 1MgCl_2_, 0.05 EGTA, 3 Mg-ATP, 0.1 cAMP, and 5 HEPES, pH 7.2 adjusted using CsOH.

During the experiments, the cells were continuously superfused with an extracellular solution containing (in mM) 138 NaCl, 5.6 KCl, 1.2 MgCl_2_, 2.6 CaCl_2_, 10 D-glucose, and 5 HEPES, pH 7.4 adjusted with NaOH at a rate of 0.4 ml/min. All electrophysiological measurements were performed at 32C. In the analysis of the measured voltage-dependent current consists of both Na^+^ and Ca^2+^ current components, were the rapid peak current (0–5 ms) represents the Na^+^ current and the sustained current during the latter part of the depolarization reflects the Ca^2+^ current. K_ATP_-currents were measured in the whole cell recording configuration, using a pipette solution (in mM) 140 KCl, 1 MgCl, 10 EGTA, 3 Mg-ATP and 10 Hepes, pH 7.2 adjusted using KOH. The cell was held at −70 mV and ± 10 mV pulses (10 ms duration) were applied alternately at a rate of 15 Hz before and after the application of 200 μM Diazoxide. In Fig.2a, membrane potential, the pipette solution contained (in mM) 76 K_2_SO_4_, 10 KCl, 1 MgCl_2_, and 5 HEPES, pH 7.3 adjusted with KOH; access was established with amphotericin (0.25 mg/mL).

### Exocytosis imaging

To visualize granule exocytosis, the cells were treated as described for electrophysiology and were additionally infected with adNPY-tdOrange2 (a well-established marker for secretory granules ^79^) and imaged after 30-36 hours using a custom-built lens-type total internal reflection (TIRF) microscope based on an AxioObserver Z1 with a ×100/1.45 objective (Carl Zeiss). Excitation was from two DPSS lasers at 491 and 561 nm (Cobolt) passed through a cleanup filter (zet405/488/561/640x, Chroma) and controlled with an acousto-optical tunable filter (AA-Opto). Excitation and emission light were separated using a beamsplitter (ZT405/488/561/640rpc, Chroma). The emission light was chromatically separated onto separate areas of an EMCCD camera (Roper QuantEM 512SC) using an image splitter (Optical Insights) with a cutoff at 565 nm (565dcxr, Chroma) and emission filters (ET525/50m and 600/50m, Chroma). Scaling was 160 nm per pixel.

Cells were imaged in a standard solution containing (in mM) 138 NaCl, 5.6 KCl, 1.2 MgCl_2_, 2.6 CaCl_2_, 10 D-glucose, 5 HEPES (pH 7.4 with NaOH). Where indicated, the GLP-1 receptor agonist exendin-4 (10 nM, Anaspec (Fremont CA)) or the K_ATP_-channel opener diazoxide (200 μM, Sigma-Aldrich) was also present. Where stated, exocytosis was evoked by elevating K^+^ to rapidly depolarize the cells (75 mM KCl equimolarly replacing NaCl in the standard solution, by computer-controlled local pressure ejection). Spontaneous glucose-dependent exocytosis (Fig.2q) was recorded for 3 min per cell after equilibration >20 min in the stated conditions.

### [Ca^2+^]_i_ imaging

SC- and primary islets were loaded with the fluorescent indicator Fura-2 LR (Ion Biosciences, San Marcos, TX, USA) by 1 h incubation with 1 μM of its acetoxymethyl ester at 37 °C in experimental buffer containing (in mM) 138 NaCl, 4.8 KCl, 1.2 MgCl_2_, 2.56 CaCl_2_, 3 D-glucose, 25 HEPES (pH set to 7.40 with NaOH), and 0.5 mg/ml BSA. After rinsing in indicator-free buffer, the islets were attached to poly-L-lysine-coated coverslips in a 50-μl chamber on the stage of an Eclipse TE2000U microscope (Nikon) and superfused with buffer at a rate of 160 μl/min. The chamber holder and 40x, 1.3-NA objective was maintained at 37 °C by custom-built thermostats. An LED light engine (LedHUB, Omicron Laserage Laserprodukte GmbH, Rodgau, Germany) equipped with 340 and 385 nm diodes and 340/26 nm (center wavelength/half-bandwidth) and 386/23 nm interference filters (Semrock, IDEX Health & Science, LLC, Rochester, NY, USA) provided excitation light that was led to the microscope via a liquid light guide. Emission was measured at 510/40 nm using a 400 nm dichroic beamsplitter and an Evolve 512 EMCCD camera (Photometrics, Tucson, AZ, USA). 340/386 nm image pairs were acquired every 2 s with the MetaFluor software (Molecular Devices Corp, Downington, PA, USA). [Ca^2+^]_i_ was calculated from the background-corrected Fura-2 LR 340/380 nm fluorescence excitation ratio from manually defined cell-sized regions of interest. The data is presented as example traces from individual islets and cells and as histograms of the [Ca^2+^]_i_ changes under different conditions normalized to the low glucose levels. All calculations were made using built-in functions of the Igor Pro 8 software (Wavemetrics, Portland, OR, USA).

### [cAMP]_m_ imaging

SC- and primary islets were transduced with adenovirus expressing the FRET-based cAMP reporter Epac-S^H188 80^ and cultured overnight. Immediately before imaging, the islets were incubated for 30 min in similar experimental buffer as for the [Ca^2+^]_i_ recordings. The islets were subsequently placed onto a polylysine-coated coverslip that was used as an exchangeable bottom of an open 50-μl chamber superfused with buffer at 200 μl/min. The chamber was mounted on the thermostated stage of a TIRF imaging setup based on an Eclipse Ti (Nikon) microscope with a 60x, 1.45-NA objective. A 445-diode laser (Cobold AB, Stockholm, Sweden) was used for excitation of the FRET donor and emission was measured at 483/32 and 542/27 (Semrock) with an EMCCD camera (DU-897, Andor Technology, Belfast, Northern Ireland, United Kingdom). The filters were mounted in a filter wheel (Sutter Instruments, Novato, CA, USA), which together with the camera was controlled by MetaFluor software. [cAMP]_m_ is expressed as the background-corrected 483/542 nm emission ratio (FRET ratio) extracted from cell-sized regions of interest.

### Metabolite tracing analysis with [U-^13^C_6_] labelled glucose

For metabolite tracing assays, 200 SC-islets or primary islets were used for each technical replicate of ^13^C_6_-glucose labelling. Islets were counted into wells of a 12-well tissue culture plate in a volume of 1ml KRB containing 3 mM unlabelled glucose. Islets were then incubated on a rotator plate at 95 RPM for 90 minutes at 37°C and 5% CO2 before being transferred to Eppendorf tubes and the basal KRB exchanged for a 0.9 ml volume of KRB containing either 3 mM (low) or 17 mM (high) [U-^13^C_6_] glucose (Cambridge Isotope Laboratories, CLM 1396). Islets were then incubated for 1 hour at 37°C and 5% CO2. After incubation, islets were washed in cold PBS before cell lysis and metabolite extraction in 75 μl of lysis buffer (80% Acetonitrile in dH2O). Islets were lysed with mild trituration before centrifugation at 10,000 g for 10 minutes at 4°C. Supernatant was transferred into Chromacol (03-FISV) MS vials with a 300 μl glass insert (Thermo-Fisher) and sealed with Chromacol caps with white pre-split septa (Thermo-Fisher), and the remaining cell pellet used for DNA quantification. Samples were analysed on a Thermo Q Exactive Focus Quadrupole Orbitrap mass spectrometer coupled with a Thermo Dionex UltiMate 3000 HPLC system (Thermo Fisher Scientific). The HPLC was equipped with a hydrophilic ZIC-pHILIC column (150 × 2.1 mm, 5 μm) with a ZIC-pHILIC guard column (20 × 2.1 mm, 5 μm, Merck Sequant). 5ul of each sample was used for each assay. Metabolite separation was achieved by applying a linear gradient of organic solvent (80–35% acetonitrile, 20 mM ammonium bicarbonate) at 0.15 ml/minute for 16 minutes at 45°C. Metabolites were analysed using heated electrospray ionization (H-ESI) with polarity switching (3400 V for positive, 3000 V for negative) at 280°C, with ion transfer at 300°C. Xcalibur 4.1.31.9 software (Thermo Scientific) was used for LC-MS control. Confirmation of metabolite peak specificity was achieved using commercially available standards (Merck, Cambridge Isotope Laboratories & Santa Cruz Biotechnology). LC-MS data quality was monitored throughout the run with running standard mixes, inhouse quality controls and blanks for detecting carry over. Peak integration and metabolite isotopologue identification was accomplished using TraceFinder 4.1 SP2 software (Thermo Scientific). Specificity of labelled peaks and isotopologues were confirmed using cell line controls, blank control samples, and non-labelled islet samples pre- and post-incubation. Natural abundance was assayed using non-labelled cell samples, and confirmed with correction calculations using IsoCor software on a subset of data^81^. To avoid any possible confounding effect of naturally occurring M+1 labelling, M+1 isotopologues were omitted from the final analyses of the relevant metabolites. Each metabolite peak area was normalized to the cell lysate DNA content or calculated as a percentage of the M+0 (non-labelled) metabolite present in the sample. DNA normalisation was also in strong agreement with levels of essential amino acids (that could not be metabolized from glucose) within each sample (data not shown). Relative abundance values are presented relative to the normalised metabolite level in primary adult islet samples in low (3 mM) [U-^13^C_6_] glucose.

### Animal experiments

Animal care and experiments were approved by National Animal Experiment Board in Finland (ESAVI/14852/2018). NOD-SCID-Gamma (NSG, Jackson Laboratories, 0055577) mice were housed in conventional facility in 12h light/dark cycle and fed standard chow. For SC-islet implantations, 250-750 SC-islets (diameter 100-200 μm), were aspirated into PE-50 tubing and compacted by centrifuging. Mice were anesthetized with isoflurane and the kidney exposed. A small opening to the kidney capsule was made and the capsule separated with a glass rod. The tubing was inserted in the opening and the SC-islets implanted using a Hamilton syringe. Kidney capsule was then closed by cautery before wound closure. Subcutaneous carprofen 5 mg/kg (Zoetis, Helsinki, Finland and buprenorphine 0.05-0.1 mg/kg, (RB Pharmaceuticals, Berkshire, UK) were used as analgesics. Non-fasted blood samples were collected from the saphenous vein monthly. Some mice received a single high-dose streptozotocin (STZ, S0130, Sigma-Aldrich) injection (130 mg/kg) 4 months after engraftment to eliminate the mouse beta cells. To verify the functionality of the SC-islet graft, the engrafted kidney was removed 1 month after the STZ injection after ligating the renal vein and artery and the ureter and removing the entire kidney. To test the functionality of the SC-islet grafts, the mice were subjected to an intraperitoneal glucose tolerance test. Mice were fasted 6-8 h before the test. The mice were weighted, and blood glucose was measured before the test. Glucose (3g/kg) was injected intraperitoneally, and blood samples (30 μl) were taken from the saphenous vein after 15 min, 30min, 60 min and 90 to measure blood glucose and circulating human C-peptide levels by ELISA (Mercodia).

### Single cell RNA sequencing sample preparation and analysis

SC-islets were incubated with 1:1 mix of TrypLe and 10x Trypsin – EDTA for 10 min in +37°C. Disassociation medium was neutralized by adding 3x volume 0.5% FBS in PBS and strained through a 30-μm filter to remove cell clusters. Finally, the cells were resuspended in PBS+0.04% BSA, washed, counted and diluted to a 1×10^6^ cells/ml concentration for encapsulation.

Single cell gene expression profiles were generated with 10x Genomics Chromium Single Cell 3’RNAseq platform using the Chromium Next GEM Single Cell 3’ Gene Expression (version 3.1 chemistry). This resulted in 220M read pairs on average per sample (with 26-28bp read 1, 8bp i7 index and 89-98bp read 2). We included two reference scRNA-seq datasets in our analysis; one sample of pancreatic progenitor cells ^43^ and 12 samples of primary adult human pancreatic islets ^44^. Raw data (fastq) processing was performed with 10x Genomics Cell Ranger (v3.1) pipeline. The reads were mapped to a hybrid of human and mouse reference genomes (GRCh38.98 and GRCm38.98). The UMI counts were filtered with DropletUtils ^82^ to remove empty droplets (FDR>=0.01), mouse cells and mouse transcripts from the data. The filtered counts were analysed with Seurat ^83^. The counts were normalized, scaled and analysed for PCA with default methods. The variable genes (top 1000) were identified separately for each sample and combined during the analysis (for a total of 6900 variable genes). To reduce biases among datasets we used Harmony ^84^ on the first 50 PCs with sample as the covariable (with theta = 2, nclust = 50, max.iter.cluster = 40, max.iter.harmony = 10). The integrated (harmonized) PCs were used to build the UMAP, find the neighboring cells (using Shared Nearest Neighbor), and identify cell clusters using default Seurat methods. To reduce background RNA contamination from disrupted cells we used SoupX ^85^ with clusters identified with Seurat, and known cell type specific marker genes (GCG, TTR, INS, IAPP, SST, GHRL, PPY, COL3A1, CPA1, CLPS, REG1A, CELA3A, CTRB1, CTRB2, PRSS2, CPA2, KRT19, VTCN1) to estimate the level of contamination. The Seurat analysis was then repeated with the adjusted counts with the following modifications. Cells with less than 1000 UMI counts or 200 expressed genes were excluded. We also removed cells with unusually high level of mitochondrial reads (>25% of counts). During clustering the resolution was adjusted to 0.2. Differentially expressed genes among clusters and sample types were identified with MAST using Seurat. The clusters were reordered by similarity and identified to cell types by the differentially expressed genes corresponding to known marker genes. We focused the analysis on the identified endocrine cells and rebuilt the UMAP and clustering on those cells. To improve gene expression representation we used a denoising and imputation method with Rmagic ^86^. We performed pseudotime analysis on the endocrine cells using monocle2 ^87^. To validate the results from the pseudotime analysis we used an independent method based on RNA velocity. The reads mapping to exons and introns were recounted with velocyto ^88^ and analysed with scVelo ^89^. Gene set enrichment analysis was performed using Enrichr^90^. Cell-cell interactions were inferred using CellPhoneDB^91^.

### Data collection and Statistical methods

Morphological data represents population-wide observations from independent differentiation experiments. Insulin secretion, respirometry and metabolomics data represents samples of independent SC-islet differentiation experiments or islet donors. Electrophysiology and measurements of Ca^2+^, cAMP and exocytosis represent recordings from individual cells in independent experiments, which are pooled depending on the differentiation experiment, or represented as individual measurements. *In vivo* data is derived from independent animals. Transcriptomics data represents data on the level of single cells, which are pooled from 2-3 independent differentiation experiments per timepoint. Statistical methods used are described in each figure legend.

## Data availability

All single-cell RNA sequencing data are deposited in the Gene Expression Omnibus -database under accession code GSE167880. All other data are available upon request from the corresponding author.

## Acknowledgements

We thank Prof. C. Wollheim for invaluable feedback on the manuscript. H. Grym, A. Laitinen and S. Eurola are thanked for expert technical support, and J. Juutila and J. Morikka for the processing and acquisition of metabolite tracing data. We thank FIMM SingleCell Analytics unit (supported by HiLIFE and Biocenter Finland) for single cell RNA sequencing services. We are grateful to the Nordic Network for Islet Transplantation (supported by the strategic grant consortium Excellence of Diabetes Research in Sweden, EXODIAB), and the IsletCore, University of Alberta, Canada, for providing human islets. This study was supported by the Academy of Finland grant 297466 and MetaStem Center of Excellence grant 312437 (to TO, VH, PK), the Sigrid Jusélius Foundation Grant (to TO), the Novo Nordisk Foundation (to TO, SB, AT), Diabetes Wellness Finland (DB, JSV) and the Helsinki University Hospital Research Funds (to TO) and an EMBO Long-Term Fellowship ALT295-2019 (to DB). Additional funding was provided by The Swedish Research Council (SB, AT, POC), Barndiabetesfonden (SB, AT, POC, JL), Swedish Diabetes Foundation (SB, AT), Family Erling-Persson Foundation (POC, TO), Family Ernfors Foundation (AT, JL), Diabetes Wellness Sweden (AT) the Innovative Medicines Initiative 2 Joint Undertaking under grant agreement 115797 (INNODIA) and 945268 (INNODIA HARVEST). This Joint Undertaking receives support from the Union’s Horizon 2020 research and innovation programme and the European Federation of Pharmaceutical Industries and Associations, JDRF, and the Leona M. and Harry B. Helmsley Charitable Trust (to TO, AT).

## Author contributions

D.B. conceived and conceptualized the study, developed differentiation protocols, differentiated SC-islets, analysed the single cell transcriptomic data and wrote the manuscript. T.B. conceptualized and performed the metabolomic analyses, differentiated SC-islets and wrote the manuscript. V.L. developed differentiation protocols, performed and analysed SC-islet differentiation-, insulin secretion- and IHC experiments and wrote the manuscript. J.S.V. developed differentiation protocols, performed and analysed differentiation and animal experiments and participated in the writing of the manuscript. O.H., O.D., P.E.L and M.Y. performed and analysed cell physiology experiments. H.M. performed and analysed differentiation and animal experiments, H.I., V.C. and J.U. participated in the differentiation experiment analysis. A.N. assisted in the single-cell transcriptomic experiments and J.K. in the data analysis. A.I.N. helped in the metabolomic analysis pipeline. H.V. and E.J. performed electron microscopical analyses. E.K., V.H. and P.K. helped in the analysis of metabolic data and participated in the manuscript writing. J.L. and P.O.C. acquired funding and participated in the differentiation and animal experiments. S. B. and A.T. supervised the cell physiology experiments, acquired funding, and participated in manuscript writing. T. O. conceived and supervised the study, provided resources, acquired funding and wrote the manuscript.

