## Supplementary_Figures for "Functional, metabolic and transcriptional maturation of stem cell derived beta cells"

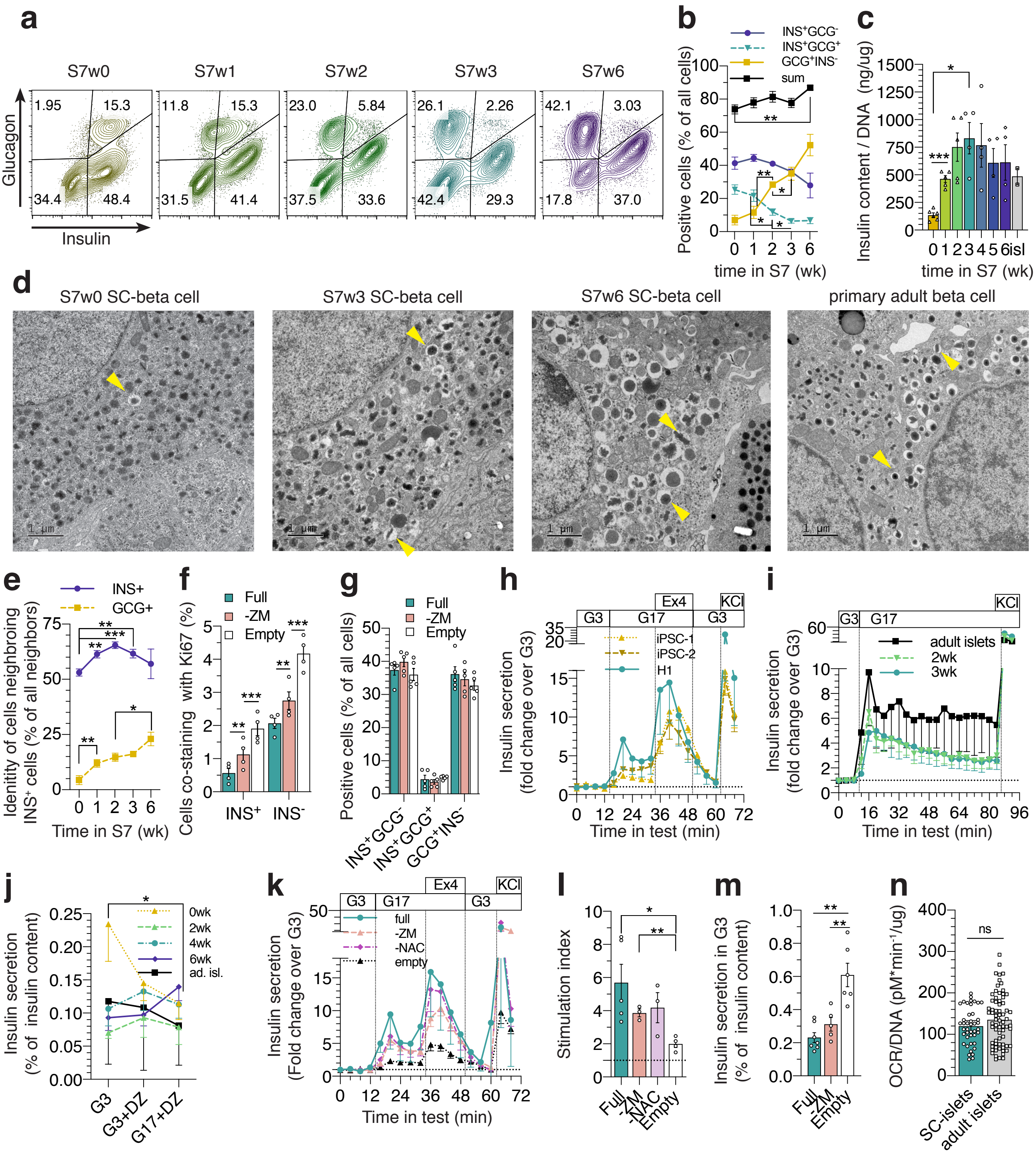
a
S7w0
S7w1
S7w2
S7w3
S7w6
Glucagon
Insulin
1.95 15.3 34.4 48.4
11.8 15.3 31.5 41.4
23.0 5.84 37.5 33.6
26.1 2.26 42.4 29.3
42.1 3.03 17.8 37.0
b
INS+GCG-
INS+GCG+
GCG+INS-
sum
Positive cells (% of all cells)
time in S7 (wk)
c
Insulin content / DNA (ng/ug)
time in S7 (wk)
isl
d
S7w0 SC-beta cell
S7w3 SC-beta cell
S7w6 SC-beta cell
primary adult beta cell
1 µm
e
INS+
GCG+
Identity of cells neighbroing INS+ cells (% of all neighbors)
Time in S7 (wk)
f
Full
-ZM
Empty
Cells co-staining with Ki67 (%)
INS+
INS-
g
Full
-ZM
Empty
Positive cells (% of all cells)
INS+GCG-
INS+GCG+
GCG+INS-
h
G3
G17
Ex4
G3
KCl
iPSC-1
iPSC-2
H1
Insulin secretion (fold change over G3)
Time in test (min)
i
G3
G17
KCl
adult islets
2wk
3wk
Insulin secretion (fold change over G3)
Time in test (min)
j
0wk
2wk
4wk
6wk
ad. isl.
Insulin secretion (% of insulin content)
G3
G3+DZ
G17+DZ
k
G3
G17
Ex4
G3
KCl
full
-ZM
-NAC
empty
Insulin secretion (Fold change over G3)
Time in test (min)
l
Stimulation index
Full
-ZM
-NAC
Empty
m
Insulin secretion in G3 (% of insulin content)
Full
-ZM
Empty
n
ns
OCR/DNA (pM*min-1/ug)
SC-islets
adult islets

**Supplementary figure 1**

**(a)** Representative flow cytometry for INS and GCG at stage 7

**(b)** Quantification of the populations in (a), n=3-7, Two-way ANOVA.

**(c)** Insulin content normalized to DNA content of the lysed SC-islets after static incubations (Fig1g), One-way ANOVA with Welch's correction

**(d)** Electron micrographs of SC-beta cells at S7 week 0,3 and 6, and of adult human beta cells. Scale bar 1 μm, yellow arrows denote mature insulin granules

**(e)** $INS^+$ and $GCG^+$ nuclei within 8 μm of $INS^+$ nuclei, percentage of all nuclei within 8 μm of $INS^+$ nuclei. Quantified from INS – GCG immunostainings. n=2-9, population-wide characterizations of individual differentiations.

**(f-g)** Quantification at S7 wk3. Comparison of the effect of different media compositions used throughout S7: ZM+NAC+T3 (Full), NAC + T3 (-ZM) and base medium without ZM, NAC or T3 (empty) quantified for **(f)** INS and Ki-67 (Matched Two-way ANOVA, n=4) and **(g)** INS and GCG, n=5.

**(h)** Insulin secretion responses of S7w3 SC-islets to stimulation from 2.8 mM to 16.8 mM glucose (G3 to G17), 50 ng/ml exendin-4 (Ex4) and 30 mM KCl in perifusion. Comparison of two wildtype iPSC lines (n=2-3) and H1 hESC-line used in the rest of the study (n=10).

**(i)** Insulin secretion response to 72-minute stimulation with G17. Normalized to secretion during G3, the first 12 minutes of the test. N=3

**(j)** Insulin secretion as %of total insulin content during 30-min incubations in G3, G3 + $K_{ATP}$-channel opener diazoxide 100 μM (DAZ) and G17 + DAZ in a static test. N=3-4 for SC-islets, N=2 for primary islets. Two-way ANOVA

**(k)** Same test as (h). Comparison of different media compositions used throughout S7: Full, -ZM, ZM + T3 (-NAC) and empty. Data points for the full media (n=5) include only data from the same differentiation batches as the data from the reduced medias (n=3-4). Normalized to the average secretion during G3

**(l)** Ratio of insulin secretion in G17 (average of 16 – 32 min) over secretion in G3 (average of 0 – 12 min) in test (k). One-way ANOVA with Welch's correction.

**(m)** Insulin secretion in G3 as percentage of total insulin content during a 30-minute incubation in a static insulin secretion assay, comparison of different media compositions used throughout S7. One-way ANOVA with Welch's correction.

**(n)** Oxygen consumption rate normalized to DNA content in G3

All data are presented as mean ± SEM. * $p < 0.05$, ** $p < 0.01$, *** $p < 0.001$

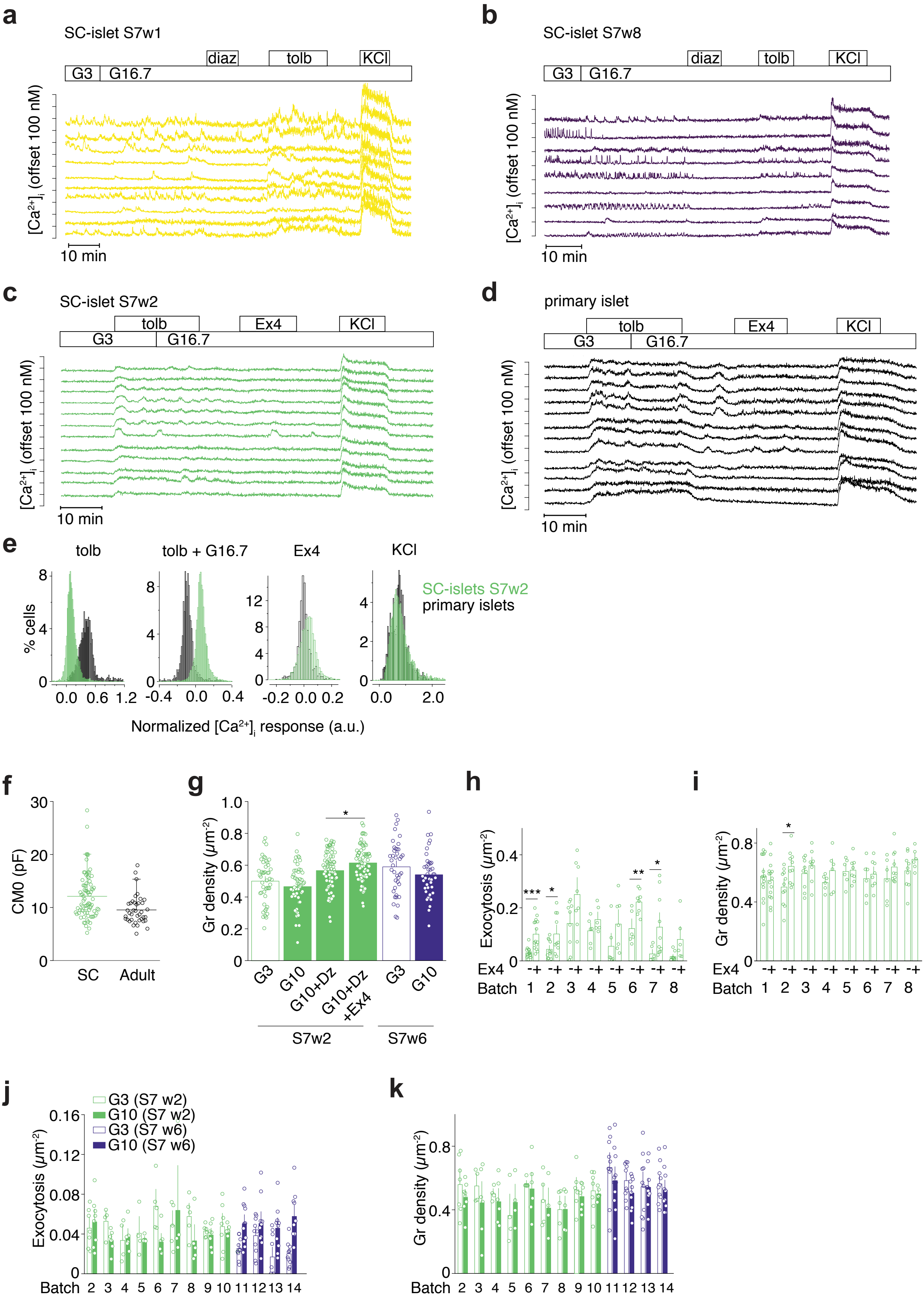

a
SC-islet S7w1
G3
G16.7
diaz
tolb
KCl
[Ca2+]i (offset 100 nM)
10 min
b
SC-islet S7w8
G3
G16.7
diaz
tolb
KCl
[Ca2+]i (offset 100 nM)
10 min
c
SC-islet S7w2
G3
tolb
G16.7
Ex4
KCl
[Ca2+]i (offset 100 nM)
10 min
d
primary islet
G3
tolb
G16.7
Ex4
KCl
[Ca2+]i (offset 100 nM)
10 min
e
tolb
tolb + G16.7
Ex4
KCl
SC-islets S7w2
primary islets
% cells
Normalized [Ca2+]i response (a.u.)
f
CM0 (pF)
SC
Adult
g
Gr density (μm-2)
G3
G10
G10+Dz
G10+Dz +Ex4
G3
G10
S7w2
S7w6
h
Exocytosis (μm-2)
Ex4
Batch
i
Gr density (μm-2)
Ex4
Batch
j
G3 (S7 w2)
G10 (S7 w2)
G3 (S7 w6)
G10 (S7 w6)
Exocytosis (μm-2)
Batch
k
Gr density (μm-2)
Batch

**Supplementary figure 2**

**(a-b)** Representative [$Ca^{2+}$]$_i$ recordings from SC-islets at S7w1 (a) and S7w8 (b) during exposure to 3 (G3) and 16.7 mM (G16.7) glucose, 250 $\mu$M diazoxide, 1 mM tolbutamide and 30 mM $K^+$.

**(c-d)** [$Ca^{2+}$]$_i$ recordings from SC-islets at S7w2 (c) and primary islets (d) exposed to 1 mM tolbutamide, G16.7, 10 nM exendin-4 and 30 mM $K^+$.

**(e)** Histograms showing the changes of [$Ca^{2+}$]$_i$ in response to various treatments from SC-islets (green; n=4079 cells) and primary islets (black, n=1187 cells).

**(f)** Whole cell capacitance as measure of cells size in SC- and primary beta cells. Symbols represent individual cells.

**(g)** Docked granules measured as average granule density, normalized to footprint area, in conditions as indicated. Dots represent individual cells. $p = 0.01$, two-tailed *t*-test

**(h)** Total exocytosis during 40 s of $K^+$-stimulation in different SC-beta cell preparations. Dots represent individual cells. Student's t-test with * for p<0.05, *** for p<0.01 and p<0.001.

**(i)** Average granule density normalized to the footprint area of different SC-beta cell preparations. Dots represent individual cells. Student's t-test with * for p<0.05.

**(j)** Total spontaneous exocytosis during 3 minutes for indicated conditions in 14 different SC-beta cell preparations. Dots represent individual cells. Student's t-test.

**(k)** Average granule density normalized to the footprint area of different batch/preparation of cells as indicated. Dots represent individual cells.

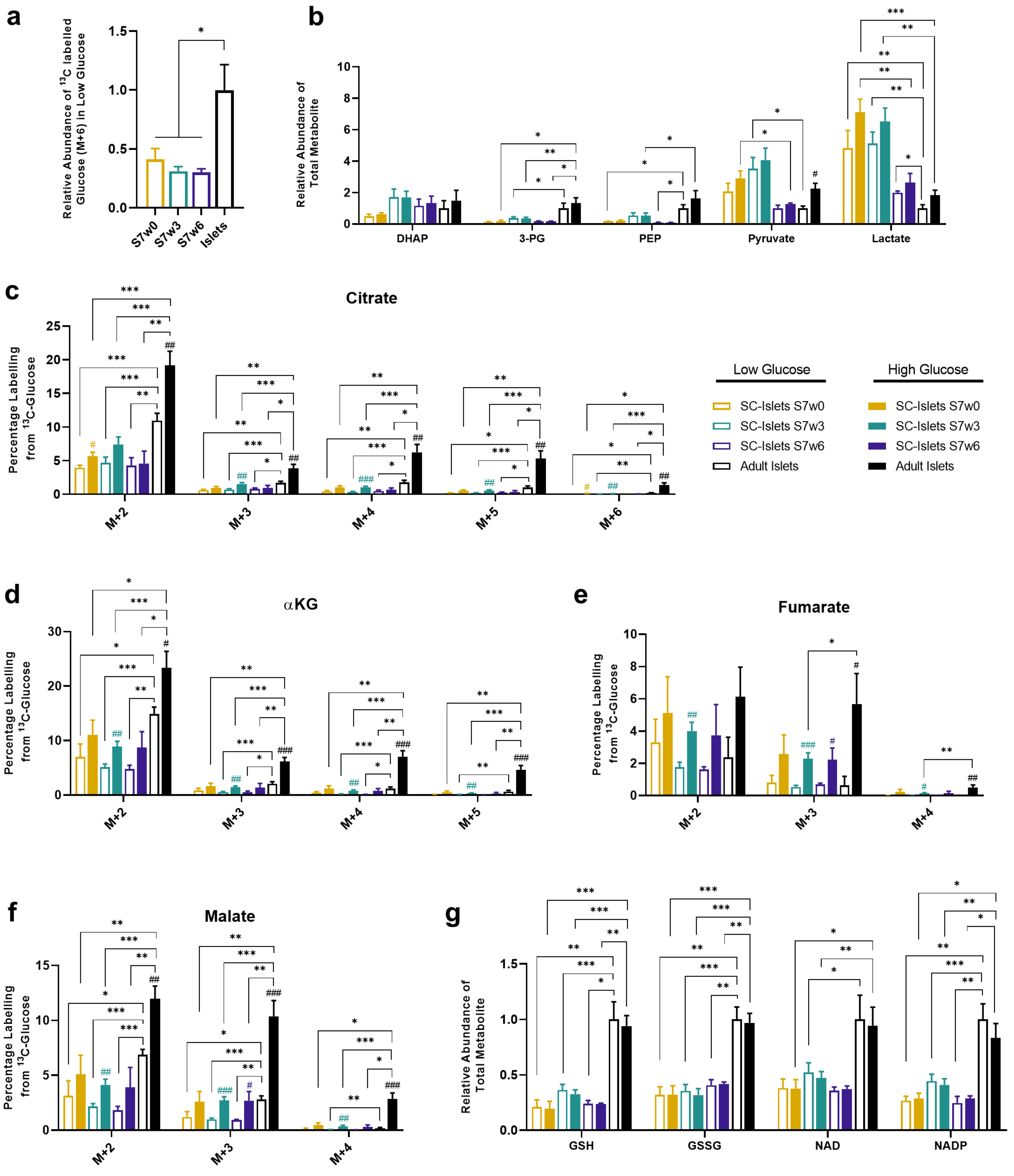
a
Relative Abundance of $^{13}$C labelled Glucose (M+6) in Low Glucose
S7w0
S7w3
S7w6
Islets
b
Relative Abundance of Total Metabolite
DHAP
3-PG
PEP
Pyruvate
Lactate
c
Citrate
Percentage Labelling from $^{13}$C-Glucose
M+2
M+3
M+4
M+5
M+6
Low Glucose
SC-Islets S7w0
SC-Islets S7w3
SC-Islets S7w6
Adult Islets
High Glucose
SC-Islets S7w0
SC-Islets S7w3
SC-Islets S7w6
Adult Islets
d
αKG
Percentage Labelling from $^{13}$C-Glucose
M+2
M+3
M+4
M+5
e
Fumarate
Percentage Labelling from $^{13}$C-Glucose
M+2
M+3
M+4
f
Malate
Percentage Labelling from $^{13}$C-Glucose
M+2
M+3
M+4
g
Relative Abundance of Total Metabolite
GSH
GSSG
NAD
NADP

**Supplementary Figure 3**

**(a)** The abundance of M+6 labelled Glucose retained under low glucose treatment of SC-islets from week 0 to week 6 of maturation, relative to adult islets.

**(b)** The total content of glycolytic metabolites (combined M+3 labelled and non-labelled) following low and high glucose exposure. All values shown relative to the content of adult islets in low glucose.

**(c-f)**, The percentage of total metabolite labelling following low and high glucose exposure for each isotopologue of the TCA metabolites citrate (c), alpha-ketoglutarate (αKG) (d), fumarate (e), and malate (f).

**(g)**, The combined total abundance of reduced and oxidized forms of glutathione (GSH and GSSG) and the reducing agents NAD and NADP in SC-islets and adult islets.

Error bars ± SEM with statistical significance determined by two-tailed t-tests. '#' symbols indicate internal significance from low to high glucose labelling, '*' symbols denote significance between SC-islet timepoints or adult islet samples at each glucose concentration. $^{\#,*}p < 0.05$, $^{\#\#,**}p < 0.01$, $^{\#\#\#,***}p < 0.001$. SC-islets S7w0 (n=4), S7w3 (n=12-13), S7w6 (n=3), adult islets (n=6).

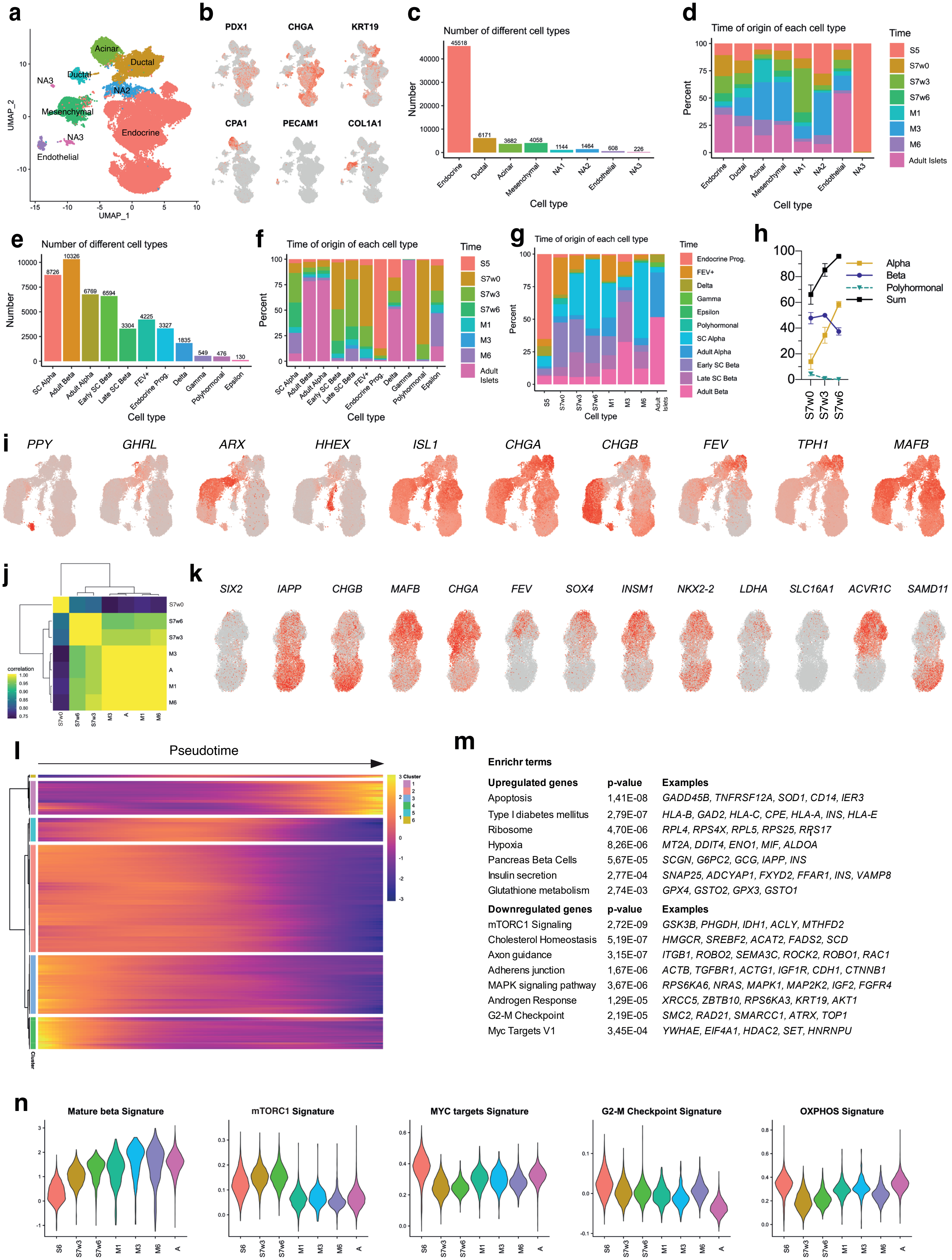
a
Acinar
Ductal
Ductal
NA3
NA2
Mesenchymal
NA3
Endocrine
Endothelial
UMAP_1
UMAP_2
b
PDX1
CHGA
KRT19
CPA1
PECAM1
COL1A1
c
Number of different cell types
45518
6171
3682
4058
1144
1464
608
226
Number
Cell type
Endocrine
Ductal
Acinar
Mesenchymal
NA1
NA2
Endothelial
NA3
d
Time of origin of each cell type
Percent
Cell type
Time
S5
S7w0
S7w3
S7w6
M1
M3
M6
Adult Islets
e
Number of different cell types
8726
10326
6769
6594
3304
4225
3327
1835
549
476
130
Number
Cell type
SC Alpha
Adult Beta
Adult Alpha
Early SC Beta
Late SC Beta
FEV+
Endocrine Prog.
Delta
Gamma
Polyhormonal
Epsilon
f
Time of origin of each cell type
Percent
Cell type
Time
S5
S7w0
S7w3
S7w6
M1
M3
M6
Adult Islets
g
Time of origin of each cell type
Percent
Cell type
Time
Endocrine Prog.
FEV+
Delta
Gamma
Epsilon
Polyhormonal
SC Alpha
Adult Alpha
Early SC Beta
Late SC Beta
Adult Beta
S5
S7w0
S7w3
S7w6
M1
M3
M6
Adult Islets
h
Alpha
Beta
Polyhormonal
Sum
S7w0
S7w3
S7w6
i
PPY
GHRL
ARX
HHEX
ISL1
CHGA
CHGB
FEV
TPH1
MAFB
j
correlation
S7w0
S7w6
S7w3
M3
A
M1
M6
k
SIX2
IAPP
CHGB
MAFB
CHGA
FEV
SOX4
INSM1
NKX2-2
LDHA
SLC16A1
ACVR1C
SAMD11
l
Pseudotime
Cluster
m
Enrichr terms
Upregulated genes | p-value | Examples
Apoptosis | 1,41E-08 | GADD45B, TNFRSF12A, SOD1, CD14, IER3
Type I diabetes mellitus | 2,79E-07 | HLA-B, GAD2, HLA-C, CPE, HLA-A, INS, HLA-E
Ribosome | 4,70E-06 | RPL4, RPS4X, RPL5, RPS25, RPS17
Hypoxia | 8,26E-06 | MT2A, DDIT4, ENO1, MIF, ALDOA
Pancreas Beta Cells | 5,67E-05 | SCGN, G6PC2, GCG, IAPP, INS
Insulin secretion | 2,77E-04 | SNAP25, ADCYAP1, FXYD2, FFAR1, INS, VAMP8
Glutathione metabolism | 2,74E-03 | GPX4, GSTO2, GPX3, GSTO1
Downregulated genes | p-value | Examples
mTORC1 Signaling | 2,72E-09 | GSK3B, PHGDH, IDH1, ACLY, MTHFD2
Cholesterol Homeostasis | 5,19E-07 | HMGCR, SREBF2, ACAT2, FADS2, SCD
Axon guidance | 3,15E-07 | ITGB1, ROBO2, SEMA3C, ROCK2, ROBO1, RAC1
Adherens junction | 1,67E-06 | ACTB, TGFBR1, ACTG1, IGF1R, CDH1, CTNNB1
MAPK signaling pathway | 3,67E-06 | RPS6KA6, NRAS, MAPK1, MAP2K2, IGF2, FGFR4
Androgen Response | 1,29E-05 | XRCC5, ZBTB10, RPS6KA3, KRT19, AKT1
G2-M Checkpoint | 2,19E-05 | SMC2, RAD21, SMARCC1, ATRX, TOP1
Myc Targets V1 | 3,45E-04 | YWHAE, EIF4A1, HDAC2, SET, HNRNPU
n
Mature beta Signature
mTORC1 Signature
MYC targets Signature
G2-M Checkpoint Signature
OXPHOS Signature
S6
S7w3
S7w6
M1
M3
M6
A

**Supplementary Figure 4**

**(a)** UMAP projection of the starting scRNAseq dataset, containing 62 871 cells, colored by different cell types.

**(b)** Relative expression of marker genes for pancreatic (*PDX1*), endocrine (*CHGA*), ductal (*KRT19*), acinar (*CPA1*), endothelial (*PECAM1*) and mesenchymal (*COL1A1*) cells.

**(c)** The number of cells from each cell type identity of the starting scRNAseq dataset

**(d)** Fractional contribution of each timepoint to each cell type identity in the starting dataset.

**(e)** The number of cells from each cell type in the endocrine scRNAseq dataset.

**(f)** Fractional contribution of each timepoint to each cell type identity in the filtered, endocrine dataset.

**(g)** Fractional contribution of each cell type identity to each timepoint in the filtered, endocrine dataset

**(h)** Percentage of cells in the alpha, beta and polyhormonal clusters and the sum of these clusters in the S7 culture timepoints.

**(i)** Relative expression of marker genes for the different endocrine cell type identities.

**(j)** Pearson correlation of the average gene expression in the cells from each different time point.

**(k)** Relative expression of beta cell marker genes in the beta cell subpopulations.

**(l)** Hierarchical clustering of the top 500 genes most differentially regulated along pseudotime (see also Supplementary Table 3).

**(m)** Gene sets enriched in the upregulated and downregulated genes along pseudotime presented in (l).

**(n)** Signatures on beta cells from each different time points of gene sets differentially regulated along pseudotime.

**a**

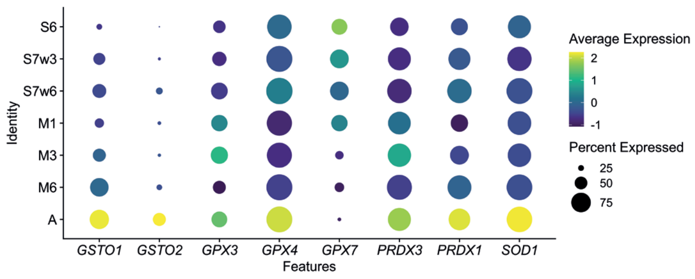


**b**

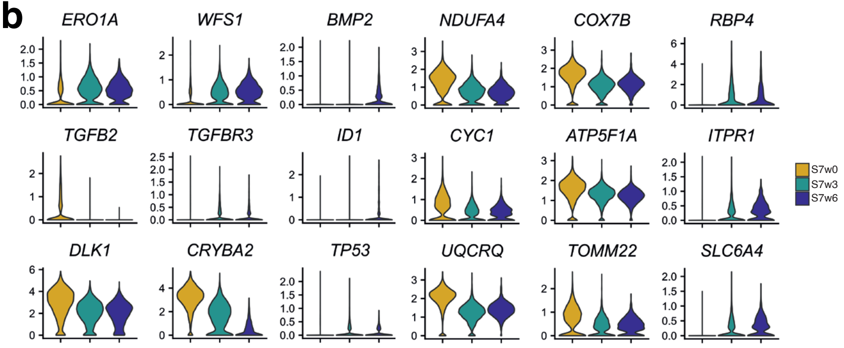


**c**

**Reactome terms**

| **S7w6 vs S7w3 SC-beta Upregulated genes** | **Raw p-value** |
|---|---|
| Respiratory electron transport (R-HSA-611105) | 3.11E-14 |
| Autophagy (R-HSA-9612973) | 7.00E-08 |
| Amino acids regulate mTORC1 (R-HSA-9639288) | 2.30E-07 |
| trans-Golgi Network Vesicle Budding (R-HSA-199992) | 7.92E-07 |
| Golgi Associated Vesicle Biogenesis (R-HSA-432722) | 5.62E-06 |
| Iron uptake and transport (R-HSA-917937) | 9.08E-06 |
| mTORC1-mediated signalling (R-HSA-166208) | 1.41E-05 |
| PERK regulates gene expression (R-HSA-381042) | 1.71E-05 |
| ATF4 activates genes in response to ER stress (R-HSA-380994) | 3.69E-05 |
| Unfolded Protein Response (UPR) (R-HSA-381119) | 3.90E-05 |
| Cristae formation (R-HSA-8949613) | 8.45E-05 |
| **S7w6 vs S7w3 SC-beta Downregulated genes** | **Raw p-value** |
| SRP-dependent protein targeting to membrane (R-HSA-1799339) | 1.88E-21 |
| Axon guidance (R-HSA-422475) | 4.73E-21 |
| Signaling by ROBO receptors (R-HSA-376176) | 1.02E-18 |
| Regulation of expression of SLITs and ROBOs (R-HSA-9010553) | 4.59E-18 |
| Formation of a pool of free 40S subunits (R-HSA-72689) | 3.85E-18 |
| Eukaryotic Translation Elongation (R-HSA-156842) | 4.22E-18 |
| Selenocysteine synthesis (R-HSA-2408557) | 3.82E-17 |
| Asparagine N-linked glycosylation (R-HSA-446203) | 2.37E-16 |
| ER to Golgi Anterograde Transport (R-HSA-199977) | 7.02E-13 |
| IRE1alpha activates chaperones (R-HSA-381070) | 2.37E-07 |
| XBP1(S) activates chaperone genes (R-HSA-381038) | 4.26E-07 |

**d**

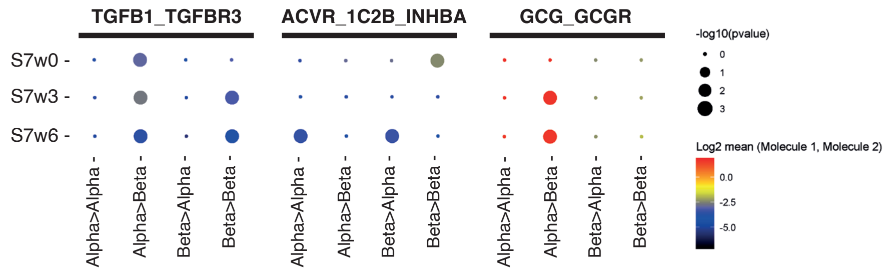

**Supplementary Figure 5**

**(a)** Expression of glutathione metabolism related genes across beta cells from different time of origin

**(b)** Expression of differentially expressed genes in the in vitro SC-beta cells.

**(c)** Reactome gene sets enriched in S7w6 upregulated and downregulated genes

**(d)** Overview of selected ligand-receptor interactions using CellPhoneDB on the in vitro SC-islet datasets. P values are indicated by circle size, the means of the average expression level of interacting molecule 1 in cell type 1 and interacting molecule 2 in cell type 2 are indicated by color, scales next to the plot.
