## Supplementary_Tables for "Functional, metabolic and transcriptional maturation of stem cell derived beta cells": Suppl table 8.pdf

**Table 1 differentiation medium composition**

| Stage | Duration (d) | Culture format | Basal medium | Additives |
|---|---|---|---|---|
| 1 | 3 | Matrigel coated planar | #1: MCDB131 (10372-019, Life Technologies)<br>+ 2 mM Glutmax ( 35050038, Life Technologies)<br>+ 1.5 g/l NaHCO3 (Sigma-Aldrich)<br>+ 5 g/l BSA fV (Sigma-Aldrich, A7030)<br>+ 5 mM glucose.<br>Total glucose 10.44 mM | + 100 ng/ml Activin A (QKine limited, UK)<br>+ CHIR (Tocris, #4423) 3.0 µM on the $1^{st}$ day, 0.3 µM on the $2^{nd}$ day, 0.0 µM on the $3^{rd}$ day. |
| 2 | 3 | Matrigel coated planar | | + 50 ng/ml FGF7 (Genscript, Z03047)<br>+ 0.25 mM Vitamin C (Sigma, A4544) |
| 3 | 2 | Matrigel coated planar | #2: MCDB131<br>+ 2 mM Glutmax<br>+ 2.5 g/l NaHCO3<br>+ 20 g/l BSA fV<br>+ 5 mM glucose<br>+ 0.5% ITS-X (supp).<br>Total glucose 10.42 mM | + 50 ng/ml FGF7<br>+ 0.25 mM Vitamin C<br>+ 0.25 µM SANT1 (Sigma, S4572)<br>+ 1 µM Retinoic Acid (Sigma, R2625)<br>+ 100 nM LDN (Selleckchem, S2618)<br>+ 200 nM TPB (Santa Cruz, Sc-204424) |
| 4 | 4 | Planar -> Aggre Well 400 | | + 10 mM Nicotinamide (N0636, SIGMA)<br>+0.25 mM Vitamin C<br>+ 100 ng/ml EGF (AF-100-15, Peprotech)<br>+ 10 µM ROCKi<br>+ 2 ng/ml FGF7<br>+ 0.25 µM SANT1<br>+ 0.1 µM Retinoic Acid<br>+ 200 nM LDN<br>+ 100 nM TPB<br>+ 10 ng/ml Activin A |
| 5 | 4 | Aggre Well 400 | #3: MCDB131<br>+ 2 mM Glutmax<br>+ 1.5 g/l NaHCO3<br>+ 20 g/l BSA fV<br>+ 15 mM glucose<br>+ 0.5% ITS-X<br>+ 10 µg/ml Heparin (H3149, Sigma-Aldrich)<br>+ 10 µM $ZnSO_4$ (Z0251, SigmaAldrich)<br>+ 1% Penicillin-Streptomycin.<br>Total glucose 20.13 mM | + 0.25 µM SANT1<br>+ 0.05 µM retinoic acid<br>+100 nM LDN<br>+ 10 µM ALK5inhII (S7233, Selleckchem)<br>+ 1 µM GC1 (4554, Tocris)<br>+ 20 ng/mL Betacellulin (100-50, Peprotech)<br>+ 100 nM GSiXX (565789, Millipore) |
| 6 | 7-8 | Suspension | | + 100 nM LDN<br>+ 10 µM ALK5inhII<br>+ 1 µM GC1<br>+ 100 nM GSiXX |
| 7 | 0-42 | Suspension | CMRL1066 (15-110-CVR, Corning)<br>+ 2 mM Glutmax<br>+ 20 g/l BSA fV<br>+ 0.5% ITS-X<br>+ 10 µg/l heparin<br>+ 10 µg/l $ZnSO_4$<br>+ 0.5 mM Sodium Pyruvate (Lonza, BE13-115E)<br>+ 1:2000 Trace elements A (25-021-CI, Cellgro)<br>+ 1:2000 Trace elements B (99-175-CI, Cellgro)<br>+ 1:2000 Lipid concentrate (11905-031, Invitrogen)<br>+ 1% Penicillin-Streptomycin.<br>Total glucose 5.32 mM | + 0.5 µM ZM447439 (ZM, Selleckchem, S1103)<br>+ 10 nM Tri-iodothyronine (T3) +<br>+ 1 mM N-Acetyl-Cysteine (NAC, A9165, Sigma-Aldrich) |
