## Supplementary_Tables for "Functional, metabolic and transcriptional maturation of stem cell derived beta cells": Suppl table 9.pdf

**Table 2 Human islet characteristics**

| Shipment# | Source | Purity (%) | Age (y) | Sex | BMI ($kg/m^2$) | HbA1c (mmol/mol) |
|---|---|---|---|---|---|---|
| 1 | Uppsala | 94 | 67 | M | 26.1 | 38 |
| 2 | Uppsala | 97 | 65 | M | 30.1 | 42 |
| 3 | Uppsala | 96 | 71 | M | 22.4 | 41 |
| 4 | Uppsala | 72 | 36 | F | 18.9 | 32 |
| 5 | Uppsala | 98 | 61 | F | 24.7 | 35 |
| 6 | Uppsala | 80 | 70 | F | 22 | 32 |
| 7 | Alberta R389 | 75 | 65 | F | 24.4 | 37 |
| 8 | Uppsala | 69 | 51 | F | 28.1 | 35 |

Uppsala = Nordic Network for Islet Transplantation, Uppsala University, Sweden
Alberta = IsletCore, University of Alberta, Canada
