## Supplementary_Tables for "Functional, metabolic and transcriptional maturation of stem cell derived beta cells": Suppl table 10.pdf

**Table 3 antibodies used**

| Epitope | Origin animal | Conjugate | Dilution | Supplier | Assay |
|---|---|---|---|---|---|
| Insulin | Rabbit | Red 647 | 1:160 | Cell Signaling Technology Cat# 9008 | FC |
| Insulin | Guinea pig | N/A | 1:500 | DAKO, A0564 | IHC |
| Glucagon | Mouse | N/A | 1:160/ 1:500 | Sigma-Aldrich, #G2654 | FC / IHC |
| Ki-67 | Rabbit | N/A | 1:500 | Leica Microsystems #NCL-Ki67p | IHC |
| Guinea pig | Goat | Red 594 | 1:500 | Thermo-Fisher, #11076 | IHC |
| Mouse | Donkey | Green 488 | 1:500 | Thermo-Fisher, #21202 | FC/IHC |
| Rabbit | Donkey | Green 488 | 1:500 | Thermo-Fisher, #21206 | IHC |

FC= Flow cytometry, IHC=Immunohistochemisty
